# An *in vivo* parallelized reporter assay to uncover tissue-specific splicing regulatory sequences in a multicellular animal

**DOI:** 10.1101/2025.08.16.670696

**Authors:** Sanjana Bhatnagar, Jade Ho, Isha Singh, Nour H. Sadek, Michael Zoberman, Bina Koterniak, Yufang Liu, Daniel Fusca, Asher D. Cutter, Alan M. Moses, John A. Calarco

**Affiliations:** Department of Cell and Systems Biology, 25 Harbord Street, Toronto, Ontario, Canada, M5S 3G5; Mayo Clinic Alix School of Medicine, Mayo Clinic College of Medicine and Science 200 1st SW, Rochester MN, 55905; Genetics and Genome Biology Program, The Hospital for Sick Children, 170 Elizabeth St, Toronto, ON M5G 1E8; Department of Ecology and Evolutionary Biology, 25 Willcocks Street, Toronto, Ontario, Canada, M5S 3B2

## Abstract

Introns play a critical role in regulating alternative splicing. However, identifying functional intronic motifs is challenging due to their short and degenerate sequence composition. Massively parallel reporter assays have provided insights into *cis*-regulatory logic governing alternative splicing, but these approaches are generally performed in cell culture, limiting their ability to capture tissue-specific contexts. Here, we implemented *in vivo* Parallelized Reporter Assays in *C. elegans* neurons and muscle cells to screen for intronic enhancer and silencer motifs among thousands of randomized sequences. We identified nearly 200 sequences regulating splicing in these tissues. We uncovered core sub-sequences with tissue-biased enhancing and silencing activity, including motifs recognized by well-characterized RNA-binding proteins, and orphan motifs with no obvious cognate binding protein. Mapping our PRA-derived motifs to native introns flanking tissue-biased alternative exons revealed their conservation across nematodes, supporting their functional relevance. Additionally, individual intronic regions frequently contained diverse combinations of these motifs, indicative of complex engagement of these sequences by RNA-binding proteins. Finally, we performed targeted mutagenesis of PRA-derived intronic enhancers flanking a neuronal microexon, identifying key *cis*-regulatory determinants of microexon splicing. Together, our study provides a framework to explore the role of intronic regions in tissue-specific splicing regulation within a multicellular organism.

**Graphical Abstract:** 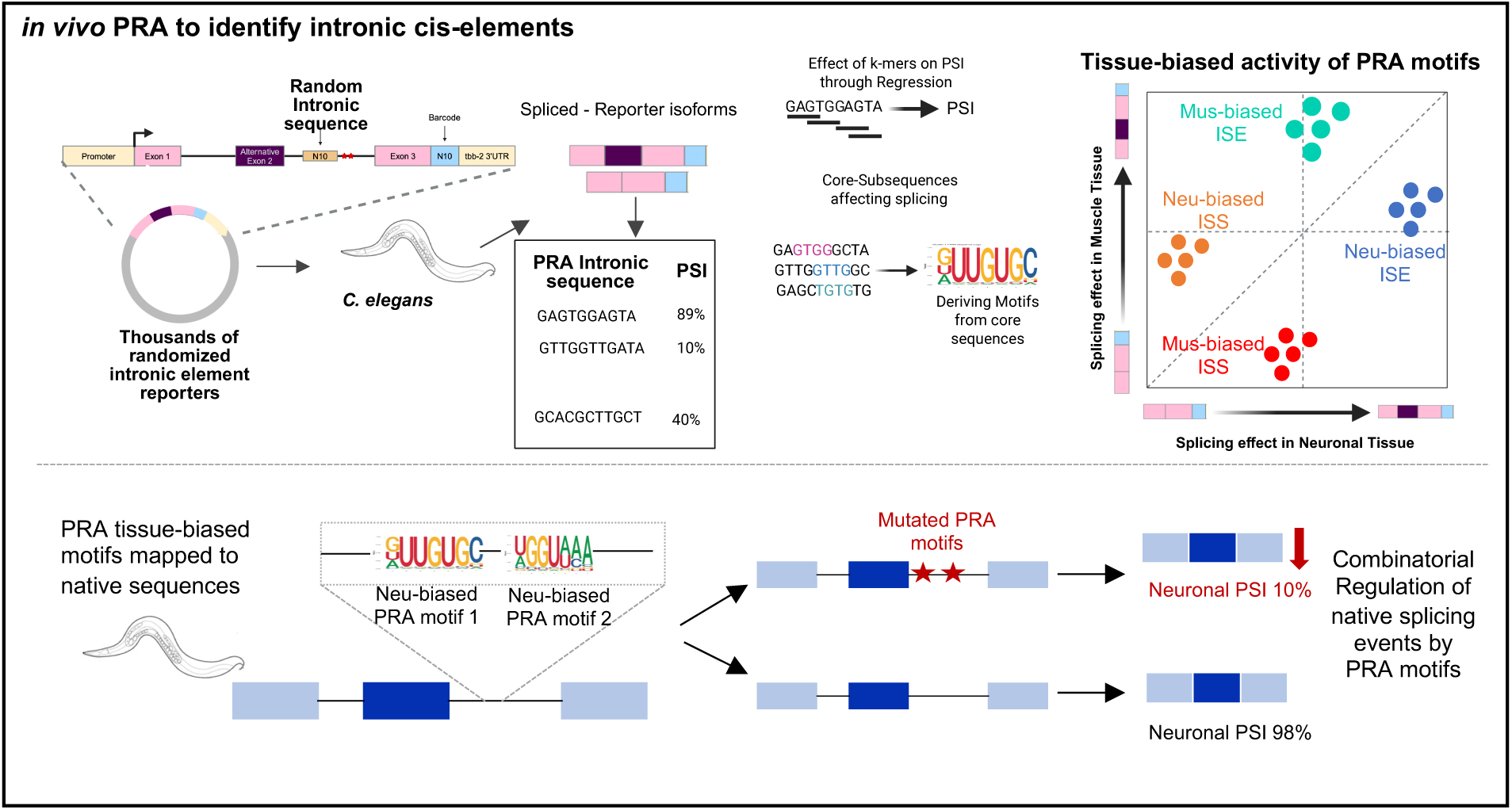

## Introduction

Pre-mRNA alternative splicing is a fundamental co-/post-transcriptional process, resulting in the removal of introns and selective joining of exons in various combinations. This mechanism vastly increases transcriptome and proteome diversity, allowing a limited number of genes to generate numerous RNA transcripts. Splice variants can confer specialized functions to widely expressed genes by being differentially expressed in a cell-, tissue-, organ-, developmental-, and physiological state-specific manner (Pan et al., 2008; Z. Wang & Burge, 2008). For example, the molecular diversity of neuronal cell types in the nervous system has been attributed, in part, to a multitude of alternative splicing events (Raj & Blencowe, 2015; S. Zheng & Black, 2013).

Intronic regions, such as the 5′ splice site, the 3′ splice site, the polypyrimidine tract, and the branch point site, form the basic framework of splicing regulation (Baralle & Baralle, 2018; Z. Wang & Burge, 2008; Warf & Berglund, 2010). In addition to these canonical elements, regulatory sequences further modulate splicing by promoting or inhibiting spliceosomal assembly at exon-intron junctions by making use of exonic and intronic splicing enhancers and silencers (Black, 2003; Blencowe, 2000; Fairbrother et al., 2002; Gao et al., 2022; Y. Wang et al., 2012). These motifs influence cell-specific splicing outcomes by recruiting RNA-binding proteins (RBPs) that are differentially expressed across cell types (for examples see: Kishore et al., 2010; Norris et al., 2014; Pedrotti et al., 2012). Therefore, characterizing these motifs in their native cellular context is crucial to understanding tissue-differential splicing regulation. Dissecting the specific effect of splicing regulatory motifs, however, remains challenging due to inherent complexity of splicing regulation. Splicing regulation depends on many factors, including multivalent *cis* elements that occur in tandem and near alternative exons (Ule & Blencowe, 2019), RNA secondary structures in nascent transcripts (Bartys, Kierzek, & Lisowiec-Wachnicka, 2019), and inputs from upstream gene regulatory layers such as transcription dynamics and chromatin states (Mikl, Hamburg, Pilpel, & Segal, 2019). These layers of complexity make it difficult to isolate the contribution of individual *cis* regulatory intronic motifs to splicing in native gene contexts.

Mini-gene splicing reporters, containing only alternative exons and their flanking intronic/exonic regions, have provided simpler sequence contexts to study the effect of *cis* regulatory elements on splicing patterns (for examples see: Cooper, 2005; Gaildrat et al., 2010; Norris et al., 2014; Rodrigues et al., 2012; Smith & Lynch, 2014; Z. Wang & Burge, 2008). For instance, early screens of randomized nucleotide sequences embedded within fluorescent reporters have successfully identified exonic splicing silencers (Z. Wang et al., 2004). More recently, deep sequencing has provided a direct readout of splicing patterns from reporters carrying libraries of randomized, rationally designed, or naturally occurring variant sequences (Baeza-Centurion et al., 2019; Ke et al., 2011; Mikl et al., 2019; Rhine et al., 2022; Rong et al., 2023; Rosenberg et al., 2015). These massively parallel reporter assays (MPRAs) have garnered immense interest for their potential to reduce regulatory complexity by testing small regions of nucleotide sequence in parallel for *cis* regulatory activity while keeping other variables unchanged.

While powerful, MPRAs have been largely limited to experiments in mammalian cell cultures, given the amenability for effective transfection and scalability. This limits the generalization of the identified *cis* regulatory elements to *in vivo* contexts and often fails to address the context-dependent regulation of biological processes, which is crucial for understanding the development of distinct tissues from the same genome. In a few recent studies, MPRAs have been implemented in organisms to study transcription factor binding sites in promoter regions and mRNA decay dynamics systematically (Agarwal et al., 2023; Kreimer et al., 2022; McAfee et al., 2022; Rabani et al., 2017). However, adaptation of MPRAs to a multicellular organism to screen for sequences with tissue-biased splicing regulatory activity *in vivo* is lacking. *C. elegans*, with an intron-rich genome and RBPs conserved across metazoa, is an attractive model system to study tissue-differential alternative splicing (Gracida et al., 2016; Koterniak et al., 2020). From a technical perspective, *C. elegans* also offers the unique capacity to accommodate large libraries of injected reporters in the form of extrachromosomal arrays (Mello, Kramer, Stinchcomb, & Ambros, 1991), making it an attractive model for *in vivo* MPRA screens.

In this study, we developed a parallelized reporter assay (PRA) to identify sequences with splicing regulatory activity in tissues of the nematode *Caenorhabditis elegans*. We used our PRA design to survey the activity of several thousand splicing reporter plasmids containing random intronic decamer sequences expressed in *C. elegans* neuronal and muscle cells. We recovered 192 intronic spicing enhancers and silencers with activity in one or both tissues. We further employed univariate linear regression to identify the core 4-7mer (k-mer) subsequences responsible for the splicing regulatory properties of these synthetic intronic elements. Our k-mer analysis identified sequences similar to known *C. elegans* RBP consensus motifs but also identified uncharacterized sequences with broad or tissue-biased splicing regulatory activity. Finally, we mapped our PRA- identified intronic silencer and enhancer sequences onto endogenous introns flanking tissue-biased alternative exons. We found that these *cis* elements are conserved and confirmed the neuronal splicing-enhancing activity of two such motifs in a model microexon splicing decision. Taken together, our results highlight the utility of our PRA approach in understanding the regulatory logic of tissue-biased alternative splicing regulation *in vivo*.

## Materials and Methods

### C. elegans strains used

We used N2 animals grown on OP50 *E. coli* for this study using standard procedures (Brenner, 1974). All animals were grown in a 21°C incubator. *Construction and sequencing of synthetic reporter libraries for parallelized reporter assay (PRA)* We utilized a previously developed *unc-16* exon-16 splicing reporter system, containing mutagenized intronic splicing enhancers (Norris et al., 2014). These enhancers are originally recognized by the trans factors UNC-75 (UGUUGUG) and EXC-7 (UAAGUU) (Norris et al., 2014). This mutagenized reporter served as a reference reporter for our experiments. Synthetic PRA element reporters were generated using forward primers containing randomized 10- nucleotide oligonucleotides, which anneal upstream of the mutated intronic *cis* elements and a reverse primer containing a 10-nucleotide barcode (IDT), annealing within the 3′ UTR. The resulting reporter fragments were then ligated using Gibson Assembly into digested vector backbones containing the pan-neuronal *rgef-1* neuron-specific promoter and the *myo-3* muscle-specific promoter and the *unc-54* 3′ UTR.

The tissue-specific reference reporter was incorporated into pools as an internal control for comparison of splicing patterns from reporters containing synthetic 10-mer PRA elements. The plasmid libraries were pooled into five groups, each containing 800 vectors per pool. The libraries were prepared for sequencing and an 8-nucleotide degenerate unique molecular identifier (UMI) downstream of barcode within the 3′ UTR using a reverse primer (5′- GAACAGTACGGTACTAAGCAAC NNNNNNNN GAAGAGTAATTGGACTCAGAAG-3′).

These samples were further purified using the MagBind Total Pure NGS (Omega Bio-Tek) protocol to enrich for fragments of the correct length, containing the synthetic PRA element, barcode, and UMI. The enriched fragments were further amplified and prepared for multiplexed plasmid sequencing using the NEBNext Ultra II DNA Library Prep Kit for Illumina. The prepared samples were assessed using Bioanalyzer and quantified using qPCR. These were then subjected to paired-end MiSeq sequencing (2x150bp). The sequencing data was further analyzed using custom Python scripts to extract true PRA element-barcode pairs using UMI counts (https://github.com/sanjanabhatnagar/in-vivoPRA-and-C.elegans-PAM-paper.git; **see README.md**). The intronic PRA elements and corresponding barcodes were extracted using the adjacent regions in the reporter. All primers and oligonucleotides used in this study are listed in **Supplementary table S1**, and plasmid maps are available upon request.

### RNA Extraction, Amplification, and Sequencing of Tissue-Specific PRA Reporter Libraries in C. elegans

The tissue-specific reporter libraries were introduced by microinjection into 𝑃_𝑜_adult animals as 50 pools, each containing 80 reporters such that targeted cells carried reporters as extrachromosomal arrays. We harvested total RNA from a mixed population of the F1 progeny of injected mothers after three days using TRI reagent (Sigma), followed by DNA digestion and cleanup using Zymo RNA Clean and Concentrator (+DNaseI) kit, according to manufacturer conditions. The RNA libraries were pooled into 5 groups, such that each pool consisted of RNA from ∼800 injected vectors. The purified RNA was reverse transcribed using Invitrogen™ SuperScript™ IV and oSB29 reverse primer (**Supplemental Table S1**), which introduced a UMI in each cDNA molecule. The resulting cDNA samples were cleaned up using 1.8X MagBind Total Pure NGS (Omega Bio-Tek) magnetic beads to cDNA ratio to remove the RT primer. We selectively enriched for reporter transcripts using Q5® High-Fidelity 2X Master Mix using primers oSB30 and oSB31 primers (**Supplemental Table S1**). This was followed by another round of cleanup using MagBind Total Pure NGS (Omega Bio-Tek) in a 1.4X beads to cDNA ratio. These samples were prepared for multiplexed sequencing using NEBNext® Ultra™ II for DNA Library Prep kit as recommended by the manufacturer. We further assessed the samples using a Bioanalyzer (Agilent) and quantified them using qPCR for accurate multiplexing before sequencing. The mixed samples were then subjected to paired end MiSeq sequencing (2 x 150bp) with 20% PhiX spike-in as internal control to improve accuracy of base calling by increasing diversity for sequencing.

### High-Throughput Classification and Quantification of Splicing Isoforms in Tissue-Specific PRA Libraries

Each tissue-specific PRA library yielded approximately 12 million reads, with a mean sequence quality (Phred score) of 34. The reads were filtered based on the quality scores and then subjected to downstream analysis. A custom Python-based splicing isoform classifier algorithm was developed to accurately categorize reads by utilizing sequence motifs such as barcodes and UMIs, and exon-exon and exon-intron junctional features (https://github.com/sanjanabhatnagar/in-vivoPRA-and-C.elegans-PAM-paper.git; **see README.md**). Initially, we extracted barcodes and UMIs from the reads by considering the flanking regions. Only UMIs that met specific length criteria (8nt) were included, while excluding a small proportion of improperly extracted UMIs and longer sequences that could result in inaccurate counts. The classifier identifies five isoform categories: included, skipped, cryptic, unspliced, and unknown. Reads are classified as spliced-in isoforms if both the Exon 1-Alternate Exon junction (TTAAAGCTAA) and the AlternateExon-Exon3 junction (TGAAAAAAGAAGATCCA) are detected. If the Exon1-Exon3 junction (TAAAGAAGAT) is present, and the AlternateExon-Exon3 junction (TGAAAAAAGAAGATCCA) is absent, the read is classified as spliced-out isoform. Furthermore, unspliced isoforms are detected based on the portions of intron1 (AATTTTTTAG) or intron2 (CAAATTTTTTCAG) present in the reads. The classifier further calculates Hamming distance for regions within each read against the abovementioned reference motifs with a mismatch threshold of less than two nucleotides. This accounts for sequencing errors while maintaining specificity and further ensures robust identification of splicing events. Reads that were not classified as spliced isoforms or unspliced isoforms due to an exceeding number of mismatches with the expected junction sequences and were thus lacking a predefined junction were categorized as unknown isoforms. This serves as a fallback category for ambiguous isoforms. Interestingly, a subset of these unknown isoforms appeared to result from cryptic splicing. We then optimized the splicing isoform classifier algorithm to classify such events as cryptic splicing isoforms. Cryptic splicing isoforms were identified by the presence of the Exon 1-Alternate Exon junction (TTAAAGCTAA) and Exon3 (AAGATCCAT), along with a sequence upstream of the inserted PRA element (GTTTCAAAT or the shorter sequence AAATTGG).

After classification of reads, UMIs were collapsed into read counts by ensuring that only unique UMI-barcode pairs were counted; duplicate pairs were removed to avoid inflation of read numbers within a category. Additionally, the reporters with more than 20 read counts were selected for downstream analysis. The percent spliced in (PSI) values were calculated using read counts of different isoforms corresponding to each PRA element and barcode. Specifically, the PSI value represents the percentage of spliced-in isoforms in the population of canonically spliced isoforms (Schafer et al., 2015).

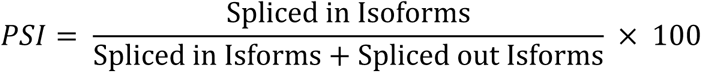

After calculation of PSI values, Fisher’s exact test was applied using spliced-in and spliced-out read counts for each PRA element reporter compared to the reference reporter. To control for false discovery rates (FDR), the Benjamini-Hochberg procedure was applied, with a threshold set at 0.01.

Similarly, cryptic isoform % represents the percentage of cryptic-splicing isoforms in the total population of spliced isoforms. To group different PRA cryptic splice site inducers, these were ranked from highest to lowest based on cryptic isoform %. We used quantile-based binning to divide the data into 5 bins containing approximately equal numbers of observations (**Fig. 2**), after sorting the elements in ascending order based on their cryptic isoform %.

### Individual Reporter Plasmid Construction and Semi-Quantitative RT-PCR validations

We prepared plasmids for a subset of selected reporters for validations and follow up experiments. In some cases, we also constructed plasmids with mutated 10-mers to confirm that ISE activity was due to the intronic sequence. Reporter fragments were amplified in two parts, the upstream portion of the reporter extending to the intronic 10-mer was amplified using TN nested F primer (for neuronal-targeted reporters) or Muscle F (for muscle-targeted reporters) along with common reverse primer unc-16 USR (**Supplemental Table S1**). The downstream portion of the reporter containing the intronic 10-mer, as well as the 10 nucleotide barcode, was amplified using a forward primer consisting the intronic PRA element and a reverse primer containing the barcode. These fragments were synthesized through PCR (Phusion polymerase, NEB) by using a neuronal *rgef-1 promoter:unc-16* minigene reference reporter plasmid or a muscle *myo-3 promoter:unc-16* minigene reference reporter plasmid as a template (**Supplemental Table S2**).

These fragments were resolved on gel and further purified for Gibson Assembly into either neuronal *unc-16* minigene vector backbone digested with SnaBI and MluI or the muscle *unc-16* minigene vector backbone digested with SnaBI and XhoI restriction enzymes. Plasmids were purified from DH5-alpha bacteria cultures using Presto Mini Plasmid kit from GeneAid. The isolated reporter plasmids were verified by Sanger sequencing (oYL 13 reverse primer) and the sequences were assessed for proper insertion of the intronic PRA elements and the barcodes (**Supplemental Table S1**).

Plasmids were delivered into 𝑃_𝑜_adult animals by microinjection. After 3 days, F1 progeny were harvested, and we used the previously stated protocol for RNA extraction and purification. This was followed by semi-quantitative RT-PCR assay using the QIAGEN OneStep RT-PCR Kit, as recommended by the manufacturer. The forward primer annealed at the start of exon 1 start (unc-16 RT F) and the reverse primer (unc-16 RT R) annealed at the end of exon 3 (**Supplemental**

**Table S1**). In general, 25-50 ng of total RNA was used as input and 30 cycles of amplification were performed. This was followed by performing 2% agarose gel electrophoresis on the obtained cDNA samples. We further performed densitometry measurements on the gel bands using Fiji ImageJ software (PSI_Densitometry_). All primer sequences used are available in **Supplemental Table S1**, and all plasmids created are available upon request.

### K-mer count Univariate Linear Regression to characterize core regulatory subsequences

To generate different k-mer count matrices, we calculated the counts of each type of k-mer (k ɛ {4,5,6,7}) in a 16 bp region including the intronic 10-mer and the flanking three nucleotides on each side. We limited the flanking length to 3bp on both sides to control for the false overrepresentation of k-mers that overlap more with the flanking region relative to the element. Each k-mer is treated as an independent predictor of the PSI value of the synthetic PRA element it belongs to. We performed univariate linear regression using the counts of only those k-mers that exist in at least five distinct synthetic PRA elements. This filter reduces noise from sparsely occurring k-mers that might not yield reliable estimates. We then performed univariate linear regression using the sklearn.linear_model package and further used scipy.stats for statistical functions in Python. We performed linear regression individually for each predictor to establish the association between the k-mer and the observed PSI value (https://github.com/sanjanabhatnagar/in-vivoPRA-and-C.elegans-PAM-paper.git; **see** README.md).

To account for measurement noise, we computed t-statistics, which inherently quantify variability by normalizing regression coefficients by their standard errors. This approach allows for a standardized assessment of the significance of each regression coefficient, facilitating comparisons across different models and datasets. The calculation of t-stat values was followed by selection of the k-mers with significantly strong effect on splicing regulation (Benjamini-Hochberg FDR, q-value = 0.05).

### Clustering tissue-biased PRA k-mers based on sequence similarity

The highly similar k-mers with significantly strong effect on splicing in both tissues were clustered using the method described in Kuret et al., 2022. The k-mers are further fragmented into subsequences and the counts of the resulting subsequences are taken across the k-mer catalogs to generate a count matrix. This represents the sequence properties of each PRA intronic k-mer and similarity is evaluated by calculating the Jaccard distance. Next, affinity propagation was performed using the same parameters as in Kuret et al., 2022, on the obtained Jaccard distance matrix to identify the appropriate number of clusters present in the data using a custom python script. This was used to guide the number of clusters defined for k-means clustering, which was subsequently performed to extract clusters of highly similar sequences (https://github.com/sanjanabhatnagar/in-vivoPRA-and-C.elegans-PAM-paper.git).

Once the k-mer clusters were obtained, they were processed to generate position weight matrices using R as previously described (Benjamin Jean-Marie Tremblay, 2014). Pairwise alignments were performed within each cluster followed by arranging them by scores (in a descending order). Then, multiple sequence alignment using ClustalW was performed. Later, position weight matrices (Benjamin Jean-Marie Tremblay, 2014), were inferred from the multiple sequence alignment for each k-mer cluster as well as consensus sequences and corresponding logos were obtained using the R package seqlogo in both tissue libraries (https://github.com/sanjanabhatnagar/in-vivoPRA-and-C.elegans-PAM-paper.git).

### Motif Comparison of Significant Splicing Enhancer and Silencer k-mers with RBP Motifs Across Tissues

We performed motif comparison analysis to compare splicing enhancing and splicing silencing k-mers against the latest *Caenorhabditis elegans* RBP motif catalogue from the RNAcompete dataset (Ray et al., 2009) using TOMTOM, a MEME suite motif tool (Howe et al., 2016). We used the Sandelin-Wasserman similarity approach to calculate motif similarity between splicing enhancing and silencing kmer cluster PWMs from both muscle and neuronal tissues by disabling the reverse complement strand search parameter. We further chose a statistical cutoff of 𝑞 − 𝑣𝑎𝑙𝑢𝑒 < 0.05 to report any significant matches.

### Phylogenetically averaged motif score (PAM) analysis on C. elegans neuronal/ muscle switch-like splicing events

In order to identify phylogenetically conserved *cis* elements by PAM score analysis, we developed a suite of scripts to extract and catalog orthologous intronic sequences based on transcript annotation files. We began by annotating all alternative and constitutive splice junctions in *C. elegans* using the coordinate and transcript information present in a recent transcript annotation file (wormbase release WBPS15, GFF and GTF files). These were classified into various splicing categories: constitutively spliced exons (always included in mRNA), alternatively spliced exons such as skipped exons (exons that are either included or excluded from the mRNA), alternative 5′ splice site (5′ss) exons (exons with two donor sites, where one is selected), alternative 3′ splice site (3′ss) exons (exons with two acceptor sites, where one is selected), retained introns (introns incorporated into mRNA), and composite exons (exons exhibiting more than one type of alternative splicing, e.g., alternative 5′ss exons that are also skipped). Additionally, since the GFF file does not include information on introns, we inferred the intron coordinates using AGAT software (Dainat, 2022). The output file contained intron coordinates inferred from exon coordinates in the GFF file. Hence, we developed a custom pipeline to filter out introns and further add intron metadata by leveraging the annotation of the exons they flanked (https://github.com/sanjanabhatnagar/Inferring-Exon-and-Intron-Metadata-from-.gff-file.git)

(Bhatnagar & Calarco, 2025). Notably, this pipeline infers splicing event metadata directly from genomic coordinates in the GFF file, without requiring expression datasets for classification. This makes it a versatile and broadly applicable tool for splicing analysis across species—applicable wherever a GFF annotation is available and not limited to the context of this study.

For this analysis, we focused on identifying orthologous flanking intron sequences from 47 switch-like splicing events differentially spliced in neuron and muscle cells (**Supplementary Table S3; B.K and J.A.C., unpublished results**) using sequences from up to 50 nematode species per gene. A minimum of five orthologous sequences per intronic fragment was required for PAMS computation to be feasible, based on orthologous groups generated in (Fusca et al., 2025) using OrthoFinder (Emms & Kelly, 2019), and the ortholog calls came from genomes obtained from WormBase ParaSite (Howe et al., 2016) and the *Caenorhabditis* Genomes Project (https://zenodo.org/records/12633738). These comprised of orthologs for the longest isoform of a given gene in *C. elegans* from the following species - *C. afra*, *C. angaria*, *C. astrocarya, C. becei, C. bovis, C. brenneri, C. briggsae, C. castelli, C. dolens, C. doughertyi, C. drosophilae, C. imperialis, C. inopinata, C. japonica, C. kamaaina, C. latens, C. macrosperma, C. monodelphis, C. nigoni, C. nouraguensis, C. oiwi, C. panamensis, C. parvicauda, C. plicata, C. portoensis, C. quiockensis, C. remanei, C. sinica, C. sp. 2, C. sp. 24, C. sp. 25, C. sp. 27, C. sp. 30, C. sp. 33, C. sp. 48, C. sp. 49, C. sp. 51, C. sp. 54, C. sp. 56, C. sp. 8, C. sulstoni, C. tribulationis, C. tropicalis, C. uteleia, C. virilis, C. waitukubuli, C. wallacei, C. yunquensis, C. zanzibari*.

We identified most similar homologs for each *C. elegans* gene by first aligning the protein sequences for all sequences per orthology group (MAFFT) (Katoh et al., 2019), followed by computing percent identity with the *C. elegans* protein and selecting the gene with encoded protein sequence with highest score per species. After identifying these groups, we selected the corresponding whole gene sequences from the species and performed whole gene sequence alignment (MAFFT). For each MAFFT run, we selected parameters --adjustdirectionaccurately to account for strandedness. This meant that all sequences will be on the same strand within an orthology group but might vary from one group to another, keeping the *C. elegans* sequence strand as reference. MAFFT adjusts the strandedness within ortholog groups as per *C. elegans* reference sequence.

Due to the unavailability of whole-genome alignments for all of these *Caenorhabditis* species, we could not leverage conventional genome liftover methods. Hence, we developed a custom alignment-based pipeline to project reference exon-intron coordinates onto orthologous sequences. This coordinate projection method employs exons as the reference in the multiple sequence alignments. The alignment approach assumes that an exon in *C. elegans* aligns to the corresponding exon in other species, and that the flanking introns align exclusively to intronic regions. Given that the alignment includes both exonic and intronic regions, the flanking intronic sequences surrounding an exon can be directly extracted and analyzed.

Consequently, the exon coordinates in the gapped sequences in alignment are deduced and further used to extract upstream and downstream intron fragments for *C. elegans* as well as the aligned orthologous sequences from other species. The length of *C. elegans* intron fragments are determined dynamically (for introns greater than 70 bp in size, a 70 bp fragment is extracted, whereas for introns shorter than 70 bp, the entire intron is considered). The intron fragment length is standardized across orthologous sequences to match the *C. elegans* intron length. This ensures that the extracted fragments from other species align with the reference species both in size and positional context. We further filtered out the orthologous sequences that are too similar (percent identity = 0.95) to the reference and ensure scores are being calculated using sequences with higher sequence divergence. This was done to avoid overestimating conservation PAM scores due to duplicated sequences between species within these groups, which may not be under evolutionary constraint but rather have not had sufficient time to diverge. Additionally, only those *C. elegans* intron fragments that had a minimum of 5 orthologous sequences were used for the analysis. In this case, all the intron fragments adjacent to the selected 47 switch exons had more than 5 orthologous sequences within the ortholog group after filtering, thus none were dropped out of the analysis. We further adopted the pipeline previously developed (Alam et al., biorxiv) to compute PAM scores for our identified tissue-biased ISE and ISS k-mer motif PWMs in intron fragment ortholog groups (https://github.com/sanjanabhatnagar/in-vivoPRA-and-C.elegans-PAM-paper.git). This resulted in a matrix with PAM scores for all PRA splicing regulatory motifs across all intron fragments. We further performed hierarchical clustering on motifs as well as intronic fragments using cluster3 (de Hoon et al., 2004) to group intronic fragments with similar motif signatures and co-occurring PRA splicing regulatory motifs in a group of sequences. The resulting clustered file was visualized using Java TreeView (Saldanha, 2004).

### Targeted mutagenesis of neuronal ISE k-mers in dpy-23 minigene reporter

We targeted neuronal ISE motifs 2 and 3 that were found to be conserved in the downstream intron of *dpy-23* microexon 4 through PAM analysis. The *dpy-23* minigene reporters with the minimal gene fragment—including the tissue-differentially spliced microexon and adjacent constitutive exons and introns—were selectively expressed in *C. elegans* neuronal tissues using the pan-neuronal promoter *rgef-1*.

The mutated *dpy-23* minigene plasmids were prepared in two parts: upstream portion was synthesized through PCR using forward primer oMZ1 and reverse primer oSB266 (Supplemental Table S6) and neuronal *rgef1:dpy-23* minigene reporter as a template. The downstream portions for three types of mutated reporters were ordered in the form of gBlocks (**Supplemental Table S1**). The *dpy-23* minigene reporter plasmid was digested using Not1 and Bsu36I restriction enzymes and Fast Digest Green buffer.

Next, we used Gibson Assembly to assemble plasmids using these fragments and the digested neuronal *dpy-23* reporter vector backbone. The assembled plasmids were purified as described above. Subsequently, we prepared microinjection mixes containing mutated neuronal *dpy-23* minigene reporters, wild-type neuronal and muscle-expressed *dpy-23* minigene reporter separately. We added pCFJ90 (*myo-2 promoter::mCherry*) at 2.5 ng/ul, and pCFJ104 (*myo-3 promoter::mCherry*) at 5 ng/uL to the microinjection mixes as co-injection markers. Microinjection mixes for each reporter were injected into 6 N2 adults each. The lines were established by picking and maintaining mCherry positive animals with stabilized extrachromosomal arrays. We then prepared 3 replicates with 4-5 plates of transgenic animals each. Each plate was seeded with 3 adults and plates were harvested after 3 days and RNA was extracted using TRI reagent (Sigma), followed by DNA digestion and cleanup using Zymo RNA Clean and Concentrator (+DNaseI) kit, according to manufacturer conditions. We further performed targeted RT-PCR, with forward primer oMZ 266 and reverse primer oMZ 175. The amplified cDNA samples were further run on 3% agarose gel for 2.5-3 hrs for high resolution of microexon spliced-in and spliced-out bands. Later, the gels from these samples were imaged and PSI densitometry values were calculated using Fiji Image J software.

## Results

*Implementation of a parallelized splicing reporter assay in C. elegans neuronal and muscle tissue* To screen for intronic elements capable of modulating splicing in different tissues, we developed a parallelized reporter assay (PRA) in *C. elegans*. Two synthetic splicing reporter libraries were constructed under the control of promoters that drive expression in either neurons or body wall muscle cells **(Fig. 1A)**. Each reporter library contained ∼4000 random 10 nucleotide (10-mer) sequences strategically located in the downstream intron flanking an alternative exon from a previously characterized *unc-16* minigene (Norris et al., 2014) **(Fig. 1A)**. Specifically, the random 10-mer sequence was positioned adjacent to a pair of mutated intronic splicing enhancers recognized by the RBPs UNC-75/CELF and EXC-7/Hu (Norris et al., 2014). We reasoned that by weakening exon inclusion in this manner, we could screen the inserted random sequences for activity from a location known to harbor *bona fide* enhancers. Each reporter was indexed with a unique 10 nucleotide barcode in the 3′ UTR, and unique molecular identifiers (UMI) were introduced in reporter transcript cDNA for de-multiplexing and precise quantification of reporter transcripts, respectively (Kivioja et al., 2012) **(Fig. 1B)**. In each library expression experiment, we also spiked in reference reporters, which lacked an inserted 10-mer sequence, to serve as a baseline for reporter splicing patterns in each tissue. This approach allowed us to measure splicing outcomes for different reporters using Percent spliced-in values (PSI), which measures the percentage of exon-included isoforms within the overall population of spliced transcripts produced by each reporter.

**Figure 1:**
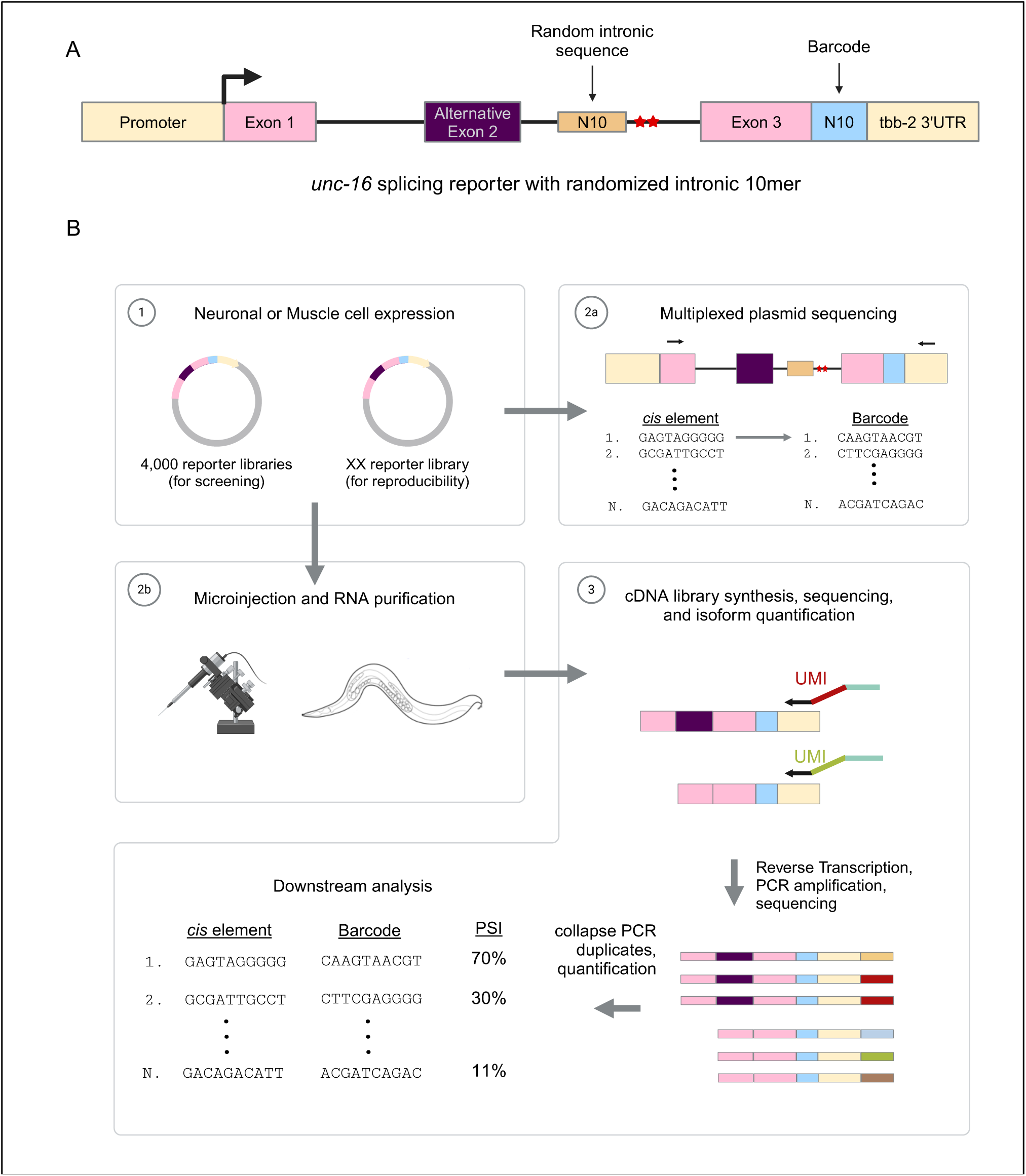
Design and Implementation of Parallel Reporter Assay (PRA) in *C. elegans* to Identify Tissue-Specific Splicing Cis-Regulatory Elements **(A)** Schematic of the minigene reporter (derived from the *unc-16* gene) used to construct tissue-specific PRA libraries. Tissue-specific expression of the reporter is achieved by introducing tissue-specific promoters (*rgef-1* promoter for neuronal tissue and *myo-3* promoter for muscle tissue-targeted libraries). Reporter includes a cassette exon (alternative exon 2), and its splicing is regulated by a random 10bp intronic element introduced in the downstream intron. Each intronic element is paired with a unique 10bp barcode located downstream of exon 3 and upstream of the tbb-2 3’ UTR. This barcode is incorporated into the reporter transcripts and enables tracking of transcripts back to their corresponding reporters. The wild-type unc-16 minigene reporter consisted of UNC-75 and EXC-7 cis-elements, which were mutated, indicated by red stars, to weaken exon inclusion. **(B)** Schematic of PRA design. (1) A library of 4000 reporters, each containing a random intronic element, is targeted to neuronal and muscle tissues separately for screening of their splicing regulatory activity. (2) A follow-up PRA library consisting of 79 reporters, targeting both neuronal and muscle tissues, was constructed as a subset of the neuronal PRA library to validate reproducibility. (2a) Multiplexed sequencing of the 4000-reporter library identifies the intronic cis-elements and barcode pairs. (2b) The 4000-reporter libraries are injected into *C. elegans* (N2 adults), and RNA is harvested and purified after 3 days. (3) The RNA samples are processed: cDNA is synthesized with unique molecular identifiers (UMIs) added during reverse transcription, followed by amplification and sequencing. Sequencing data is analyzed, and isoforms originating from each reporter (identified using barcodes) are quantified using UMI counts, eliminating PCR bias. Percent spliced-in values (PSI) are calculated for each intronic cis-element and barcode pair.

**Figure 2:**
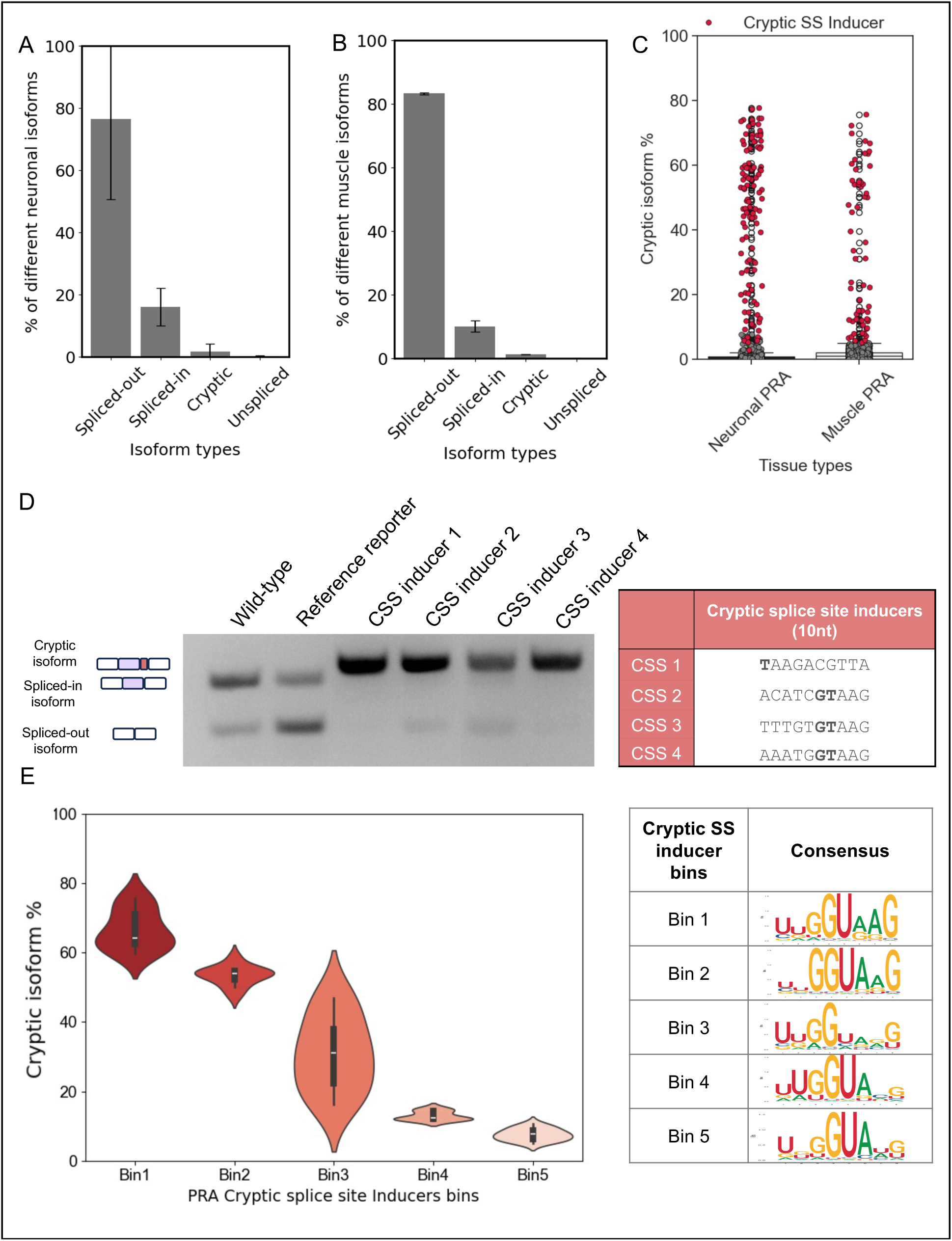
Downstream Data Analysis of PRA RNA-seq Libraries Reveals Dominance of Spliced Isoforms and Cryptic Splice Site Induction by PRA Elements **(A, B)** Bar plots showing the distribution of different isoform types observed in neuronal and muscle tissue after sequencing. The majority of the reads correspond to spliced isoforms. **(C)** Analysis of non-canonical isoforms reveals the presence of cryptic splice site (CSS) inducing PRA elements. These elements were quantified for each PRA cis-element and barcode pair in both tissue libraries. Many elements acting as cryptic splice-site inducers produce cryptic isoforms to varying extents, with cryptic isoform percentages reaching up to 80%. **(D)** Four cryptic splice site inducers identified from RNA-seq data analysis, and neuronal *unc-16* wild-type and reference reporters as controls, were validated using a semi-quantitative RT-PCR approach. The sequences of these cryptic splice site inducers are present in the adjacent table. **(E)** Binning of the cryptic splice-site inducers based on their cryptic isoform percentages reveals distinct patterns. PWMs and consensus sequences were generated for each bin. Elements in bin 1, with the highest cryptic isoform percentages, closely resemble the 5’ splice site consensus motif, while elements in bin 5 show a weaker match to the motif.

After injecting each library into animals, purifying RNA, and sequencing cDNA libraries, we successfully recovered isoform reads for 1299 and 3526 reporters from the neuron- and muscle- expressed libraries, respectively **(Supplemental Fig. S1A; Supplemental Table S4 and S5 and see methods)**. We confirmed that each nucleotide position of the inserted 10-mers was represented by all four bases without any systematic bias **(Supplemental Fig. S1B)**. To demonstrate reproducibility of our PRA results, two biological replicates of a smaller-scale PRA were also generated using a subset of 71 randomly selected reporters from the larger neuronal tissue reporter library (**Supplemental Table S6**). A strong correlation was observed between replicate PSI values from the 71 reporters when they were expressed in neurons or in muscle cells (𝑅^2^: 0.98, 𝑝: 6.64𝐸 − 49) **(Supplemental Fig. S1C and S1D)**. PSI value measurements were also positively correlated between the 71-reporter library and original 4000-neuronal reporter library (𝑅^2^: 0.61, 𝑝: 3.56𝐸 − 11) **(Supplemental Fig. S1E)**. Taken together, our results indicate that *C. elegans* can be used to survey thousands of reporters in parallel to monitor the impact of introduced random sequences on alternative splicing patterns *in vivo*.

### PRA sequences resembling a strong 5′ splice site consensus lead to cryptic alternative splice site usage

Alternative splicing occurs through the relative ability of the spliceosome to recognize competing splice sites (Aldalaqan et al., 2022; Sibley et al., 2016). One determinant of splice site usage is the relative strength of the 5′ splice sites, generally dictated by their complementarity to the first nine bases of U1 snRNA (Rinke et al., 1984; Roca et al., 2013). Prior PRA experiments have noted that random sequences inserted into reporters can occasionally trigger cryptic splice site usage (Mikl et al., 2019.; Perchlik et al., 2024). Given the location of our inserted 10-mer sequences downstream of the native 5′ splice site flanking the reporter alternative exon, we reasoned that some 10-mer sequences could similarly generate cryptic splice sites in our reporter pre-mRNA.

To characterize this possibility, we searched our reporter library sequencing data for cDNAs with unexpected exon junction connectivity (**Fig. 2**). The neuronal- and muscle-expressed reporter transcript populations were primarily composed of canonical spliced transcripts (90% and 93.22% respectively; **Fig. 2A and 2B**). However, we also identified non-canonical splice-isoforms, representing cryptic splicing events at novel intronic 5′ splice sites introduced by a subset of synthetic 10-mer elements (**Fig. 2A and 2B**). The overall prevalence of such transcripts was limited to 1.95% of all neuronal transcripts and 1.21% of all muscle transcripts **(Fig. 2A and 2B)**.

At a single reporter level, we identified a total of 54 10-mer sequences generating cryptic isoforms at a rate of >=5% (**Fig. 2C and Supplementary Table S7**) from the neuron- and muscle-expressed libraries. We independently verified cryptic splice site usage resulting from four of the top cryptic splice sites inducing reporters by semi-quantitative RT-PCR, consistent with our analysis of the sequencing data (**Fig. 2D**). To assess sequence characteristics of cryptic 5′ splice sites of varying strengths, we grouped the cryptic splice site inducing reporters into five bins based on cryptic isoform usage, ranging from 5% to 80% **(Fig. 2E)**. The sequences in each bin were aligned at the 5′ splice site to generate position weight matrices (PWMs) and consensus sequences. As expected, the bin containing reporters with the highest cryptic isoform usage rates (59.7%- 75.6%) most closely matched the 5′ splice site consensus (C/A)AG|GUAAGU-3′ (Buratti et al., 2007). Moreover, bins with lower cryptic isoform usage rates displayed progressively weaker matches to the consensus 5′ splice site sequence **(Fig. 2E)**. Interestingly, in Bin 3, sequences enabling moderate levels of cryptic splice site usage included non-canonical nucleotides such as a C at intronic position +2 (**Fig. 2E and Supplemental Table S7**). Collectively, these results demonstrate that sequences resembling consensus 5′ splice sites are sufficient to compete with weak alternative splice sites for recruitment of the splicing machinery. Our results further indicate that our PRA can recover sequence features dictating cryptic splice site usage.

### Synthetic PRA sequences can act as intronic splicing enhancers or silencers

We next determined whether our PRA approach could identify intronic 10-mers in our libraries that significantly altered PSI values when compared to a reference reporter control (|ΔPSI| = X𝑃𝑆𝐼^𝑃𝑅𝐴^𝑡𝑖𝑠𝑠𝑢𝑒 − 𝑃𝑆𝐼^𝑅𝑒𝑓𝑒𝑟𝑒𝑛𝑐𝑒^𝑡𝑖𝑠𝑠𝑢𝑒 X; |ΔPSI| > 10%; p < 0.05, Fisher’s Exact Test followed by Benjamini-Hochberg FDR critical value < 0.01). We also required that any candidate regulatory 10-mers led to minimal (<5%) cryptic splicing in reporter transcripts. Importantly, using these criteria, we recovered fifty-six 10-mers with putative intronic splicing enhancer (ISE) activity, and twenty-nine 10-mers acting as putative intronic splicing silencers (ISSs) from our neuronal-expressed reporter library (Supplemental Table S1). From our muscle-expressed reporter library, we recovered seventy-three 10-mers with putative ISE activity, and two 10-mers acting as potential ISSs (**Fig. 3A**; supplemental Table S2). To assess the robustness of the PRA in recovering ISEs and ISSs, we independently measured the splicing patterns of reporters containing the top ten 10- mers with ISE activity from the neuron-expressed library by semi-quantitative RT-PCR and densitometry **(Fig. 3B)**. All the reporters tested displayed increased alternative exon inclusion compared to the reference reporter control and generally correlated with PSI values measured by our PRA **(Fig. 3B)**.

**Figure 3:**
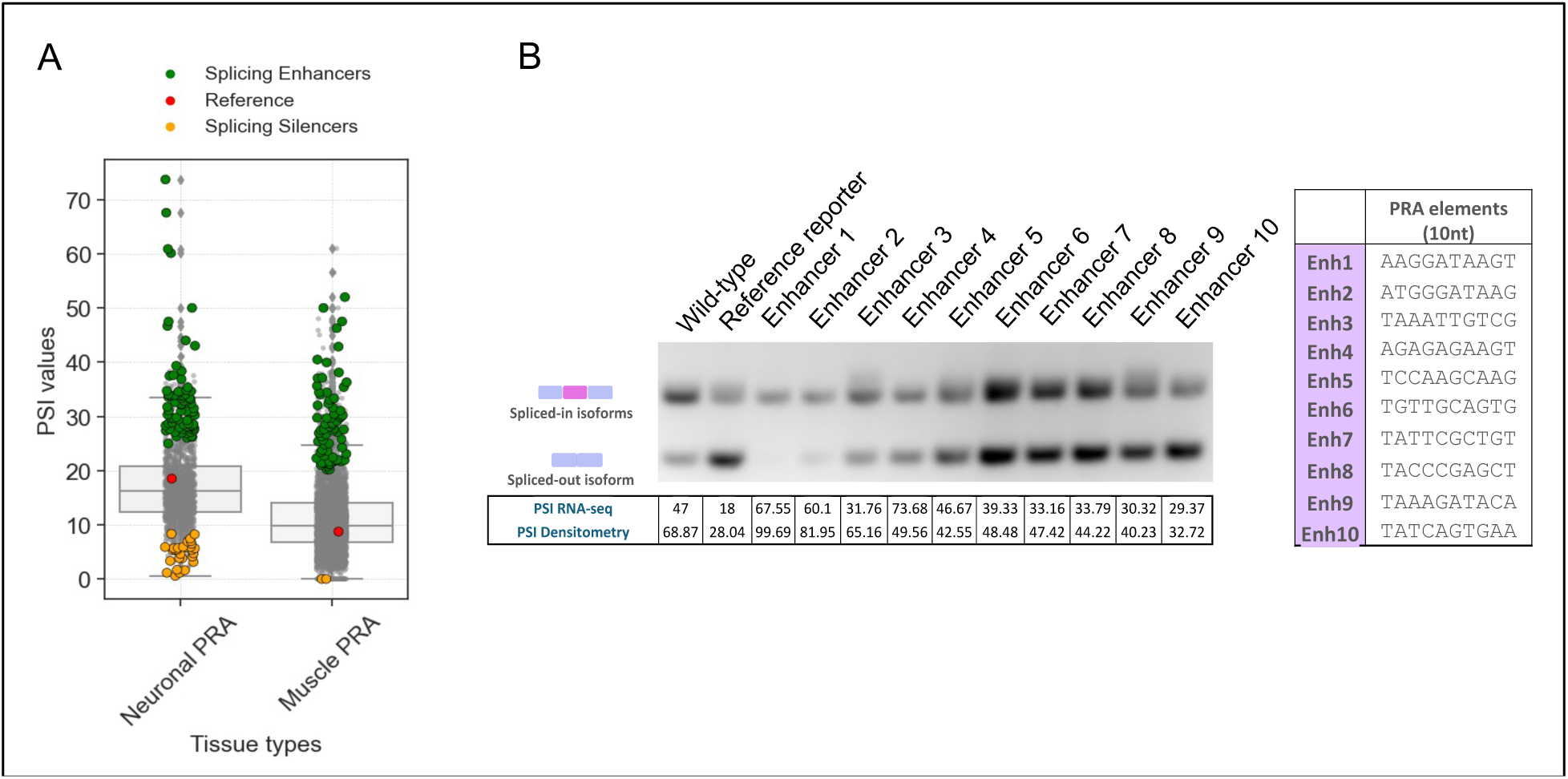
Identification of tissue-specific intronic splicing enhancers (ISEs) and intronic splicing silencers (ISSs) **(A)** Sequencing data analysis resulted in the identification of tissue-specific PRA cis-elements with PSI values significantly higher than the reference reporter (PRA ISEs) and with PSI values significantly lower than the reference reporter (PRA ISSs). (X𝑃𝑆𝐼^𝑃𝑅𝐴^𝑡𝑖𝑠𝑠𝑢𝑒 − 𝑃𝑆𝐼^𝑅𝑒𝑓^𝑡𝑖𝑠𝑠𝑢𝑒 X = ΔPSI; ΔPSI > 10%; p<0.05, Fisher’s Exact Test followed by Benjamini-Hochberg FDR critical value < 0.01). **(B)** Ten neuronal PRA ISEs were selected for semi-quantitative RT-PCR validation, and gel densitometry-based PSI values (PSI_densitometry_) are displayed along with RNA-Seq PSI values. The sequences of these PRA ISEs are presented in the accompanying table.

To test whether the observed splicing regulatory effects above can be confidently attributed to the introduced intronic 10-mers versus the ten nucleotide barcodes located in the reporter 3′ UTRs, we arbitrarily and minimally mutated the intronic sequences of 5 reporters with neuronal PRA ISEs **(Supplemental Fig. S2)**. In all cases tested, we observed a reduction in exon inclusion in reporters with mutated 10-mers compared with reporters containing the putative enhancer sequences **(Supplemental Fig. S2)**. Taken together, our results indicate that our PRAs can recover intronic sequences with splicing regulatory activity *in vivo*.

### Regulatory activity of recovered PRA cis elements can be attributed to core subsequences

Motifs associated with alternative splicing regulation *in vivo* are generally 4-7 nucleotides long (Liu et al., 1998; M. Chen & Manley, 2009; Y. Lee & Rio, 2015). Therefore, we tested shorter subsequences within our library of 10-mer intronic elements for their regulatory potential (**Fig. 4**) and assessed if these short sequences were associated with tissue-biased splicing activity (**Fig. 5 below**). We used k-mer count univariate linear regression (ULR) to comprehensively capture the contribution of 4-7 nucleotide subsequences to observed splicing outcomes. K-mer count matrices were generated for the 16 bp region containing the randomized 10 bp synthetic PRA sequence and 3bp of flanking intronic region on both sides to account for regulatory motifs created due to adjacent sequences in the reporter **(Fig. 4A)**. Based on the diversity of our PRA libraries, 100% of all possible 4-mers and, 87.8% and 99.7% of all possible 5-mers in neuronal and muscle libraries, respectively, were tested **(Fig. 4B)**. We used the t-stat value as a measure to quantify the effect of each k-mer on the splicing outcome as it inherently accounts for the uncertainty in the regression coefficient estimates by incorporating standard error **(Fig. 4C-4D; see methods for details)**.

**Figure 4:**
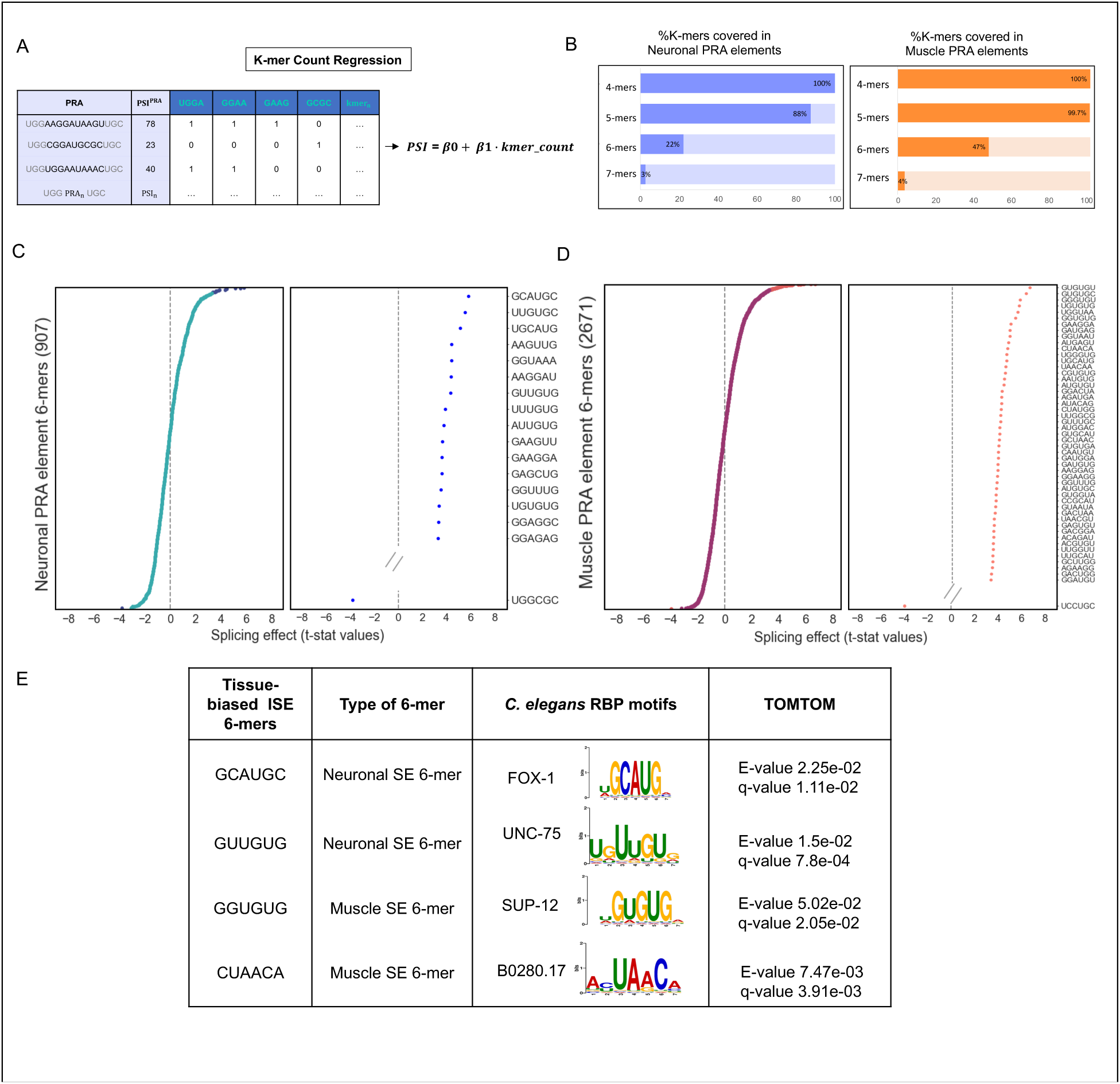
k-mer Count Regression Analysis results in identification of core Regulatory Determinants of Splicing in PRA Elements. **(A)** Schematic showing K-mer Count Regression analysis workflow. The PSI values for these PRA elements were then regressed over these k-mer counts and we obtained regression weights, symbolizing the magnitude of effect that k-mer has on splicing regulation. **(B)** The bar plot shows that all the 4-mers and a major fraction of 5-mers were covered in both neuronal and muscle tissue PRA libraries. **(C and D)** Splicing effects of 907 neuronal 6-mers and 2671 muscle 6-mers are reported by t-stat values (neuronal – C; muscle – D). The t-stat values for all recovered 6-mers are reported (X-axis) and a subset of neuronal and muscle PRA element 6-mers with significantly strong splicing effects are visualized on the right panel of each plot (Y-axis) (Benjamini-Hochberg, FDR < 0.05). **(E)** Motif comparison analysis performed using TOMTOM revealed the similarity between certain tissue-biased PRA SE (Splicing Enhancing) 6-mers and known RBP motifs such as UNC-75, FOX-1, SUP-12 and B0280.17.

**Figure 5:**
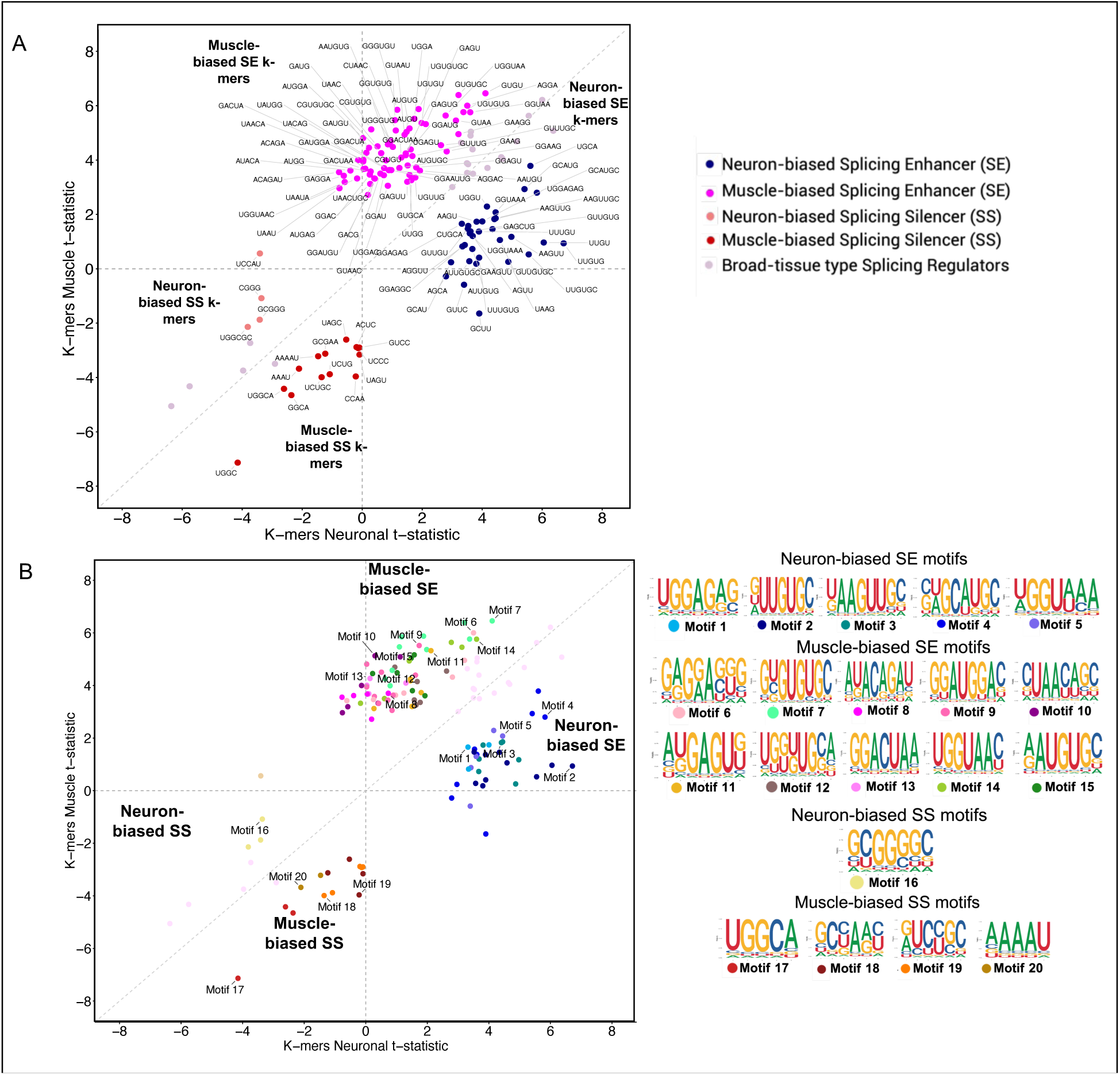
Direct comparison of tissue-specific ISE and ISS kmers reveals biases in nucleotide arrangements and potential matches to some tissue-specific RBPs. **(A)** PRA-ISE and ISS k-mers common to neuronal and muscle tissue k-mer catalogues were analyzed for tissue-biased splicing effects by comparing their t-stat values in both tissues. Four types of classes capturing tissue-specific activity of k-mers are: Neuron-biased splicing enhancing (SE) k-mers with a significantly higher t-stat value in neuronal tissue vs. muscle tissue; Muscle- biased SE k-mers with significantly higher t-stat value in muscle tissue vs. neuronal tissue; Neuron-biased splicing silencing (SS) k-mers with a significantly lower t-stat value in neuronal tissue vs. muscle tissue; and Muscle-biased SS k-mers with significantly lower t-stat value in muscle tissue vs. neuronal. **(B)** Tissue-biased k-mer cluster motifs were generated based on sequence similarity. In total, 5 neuron-biased splicing enhancing k-mer motifs (NeuSE Motif 1 – Motif 5), 10 muscle-biased splicing enhancing k-mer cluster motifs (MusSE Motif 6 – Motif 15), 1 neuron-biased splicing silencing k-mer cluster motif (NeuSS Motif 16) and 4 muscle-biased splicing silencing k-mer cluster motifs (Mus SS Motif 17 – Motif 20) were identified. The k-mers in each cluster are represented by cluster-specific colored data points in the cartesian plot and the corresponding motif logos are shown.

Using this approach, a total of 72 and 185 intronic PRA element k-mers with significant splicing regulatory effects were identified from our neuron- and muscle-expressed reporter libraries, respectively (two-tailed t-test, p-value <0.05; Benjamini-Hochberg FDR < 0.05) **(Figure 4C and 4D; supplemental Table S8 & S9)**. Here, we focus on reporting features of 6-mers, as most identified 4-mers and 5-mers could be mapped within a 6-mer context (**Supplemental Table S8 and S9, and Fig. 5 below**). We assessed whether these 6-mers could be mapped to experimentally characterized *C. elegans* RBP motifs using TOMTOM (statistical cutoff of q-value<0.05) (S. Gupta et al., 2007) with *C. elegans* RBP PWMs derived from RNAcompete datasets (**Fig. 4E**) (Ray et al. 2013). Interestingly, we recovered several 6-mers bearing similarity to known RBP consensus motifs. From our library expressed in neurons, the 6-mer with the strongest ISE activity was GCAUGC (t-stat = 5.82), followed by UUGUGC (t-stat = 5.55) **(Fig. 4C)**. These sequences bear a strong resemblance to the consensus sequences of FOX-1/ASD-1 (Jin et al., 2003; Sun et al., 2012) and UNC-75 (Kuroyanagi et al., 2013; Norris et al., 2014; L. Chen et al., 2016), known RBP splicing regulators expressed in neurons (**Fig. 4E**). From our muscle-expressed library, the 6-mer with the strongest ISE activity was GUGUGU (t-stat = 6.9), which closely resembles the consensus sequence recognized by SUP-12, a known muscle-expressed RBP splicing regulator (Anyanful et al., 2004; Kuwasako et al., 2014) **(Fig. 4D and 4E).**

In addition to recovering sequences resembling characterized RBP consensus motifs, our k-mer analysis also identified sequences with putative ISE and ISS activity not presently mapped to a cognate *C. elegans* RBP. For example, the most potent splicing-silencing 6-mers, UGGCGC (neuron-expressed library; t-stat = -3.81) and UCCUGC (muscle-expressed library; t-stat = -3.98) shared no resemblance with any known RBP motifs **(Fig. 4C-4D)**. GA- rich 6-mers with ISE activity were also identified from both tissue-expressed libraries. Additionally, AAGGAU (neuron-expressed library t-stat = 4.37) and GAAGGA (muscle-expressed library t-stat = 5.04) were identified, however, did not match any of the available *C. elegans* RBP motifs (Fig. 4C). Collectively, our results demonstrate that our PRA can facilitate the identification of both known and novel *cis* elements with splicing regulatory activity.

### Direct comparison of intronic enhancer and silencer activity in neuronal and muscle tissue reveals tissue-biased roles for sequences

Tissue-biased splicing patterns are generated through the cell-specific activity of RBPs binding to cognate *cis* elements (Fu & Ares, 2014; Feng et al., 2021; Yang et al., biorxiv). As such, we conducted a direct comparative analysis of the identified k-mers to investigate the differential effects of sequences on tissue-specific splicing regulatory activity, using the tissue-specific t-stat as a metric **(Fig. 5; see methods for details)**. In total, 21 k-mers exhibited similar splicing effects in both tissues and were classified as broad-tissue type splicing regulators **(Fig. 5A; Supplemental Table S10)**. In contrast, 119 k-mers (4-7 nucleotides) displayed tissue-biased splicing effects, with 82 k-mers (68 enhancing; 15 silencing) exhibiting muscle-biased and 37 k-mers (33 enhancing; 4 silencing) exhibiting neuron-biased splicing effects **(Fig. 5A; Supplemental Table S10)**.

As mentioned above, a subset of smaller k-mers with tissue-biased activity were subsequences of larger 6-mers and 7-mers. To further group the sequences, k-mers in each tissue-biased regulatory category were clustered based on sequence similarity using a previously described k-mer clustering and alignment method (Kuret et al., 2022; see methods for details). This approach yielded 20 intronic splicing regulatory motif clusters, including 5 neuron-biased ISE clusters, 10 muscle-biased ISE clusters, 1 neuron-biased ISS cluster, and 4 muscle-biased ISS clusters **(Fig. 5B; Supplemental Table S10)**. We assessed whether these tissue-biased PRA k-mer motif PWMs could be mapped to experimentally characterized *C. elegans* RBP motifs **(Supplemental Fig. S3).** Consistent with our earlier analysis, and with recovering *bona fide* tissue-biased splicing regulatory elements, 5 PRA k-mer motifs were matched with presently known *C. elegans* RBP splicing regulator motifs. Neuron-biased ISE motif 2 (DUUGUG-) resembled the UNC-75 consensus sequence, while motif 4 (--GCAUG-) closely matched the FOX1/ASD-1 consensus motif (**Supplemental Fig. S3**). Similarly, muscle-biased ISE motif 7 (- YGUGUG-) mapped to the consensus sequence recognized by SUP-12 and motif 10 (-UAAYW-) mapped to B0280.17 or K07H8.9 (homologs of the mammalian QKI splicing factor) motifs (**Supplemental Fig. S3**).

Interestingly, the other 16 PRA k-mer motifs with significant tissue-biased splicing regulatory effects did not map to any motifs within the *C. elegans* motif catalog (**Supplemental Table S10**). This latter group of *cis* elements represent uncharacterized motifs that warrant further study for their role in splicing regulation. Collectively, these results indicate that our PRA data can be used as a discovery tool to extract meaningful *cis* regulatory signals with tissue-biased activity.

### PRA-derived ISE and ISS motifs can be mapped to endogenous conserved intronic regions flanking tissue-biased switch-like exons

Given that our PRA-derived ISE and ISS k-mer motifs were recovered from engineered splicing reporters, we wanted to determine whether they are also functional in native sequence contexts. Specifically, we wanted to assess if the identified tissue-specific ISE and ISS sequences are conserved in intronic regions flanking tissue-biased alternative exons of *Caenorhabditis* genes. However, non-coding regions such as introns are highly diverged and can be difficult to align with traditional alignment software. For instance, several studies have demonstrated that *Caenorhabditis* nematodes undergo extensive turnover of introns (Roy & Penny, 2006; Ma et al., 2022; Teterina et al., 2025). However, prior work has demonstrated that intronic regions flanking alternative exons tend to display elevated levels of sequence conservation compared to regions flanking constitutive exons, likely reflecting selective pressures acting to retain splicing regulatory sequences (Sugnet et al., 2004; Baek & Green, 2005; Fagnani et al., 2007; Koterniak et al., 2020).

To measure conservation of specific ISE and ISS motifs, we used phylogenetic average motif (PAM) score representations for intronic fragments (Alam et al., biorxiv). This approach averages the best match scores for a provided motif (represented as a position weight matrix) across sets of unaligned orthologous non-coding regions. The scores are further normalized by scrambling the motifs 100 times and calculating a z-score between the real match scores and the distribution of scrambled match scores serving as a background (Alam et al., biorxiv). (**see Methods for further details**). These z-scores are hereafter referred as PAM scores. An advantage of this approach is that it leverages information about conservation of motifs while also accommodating some flexibility in their positioning within a non-coding region.

Mining existing transcriptome data from *C. elegans* tissues, we have identified a set of 47 alternative exons in genes that display switch-like splicing patterns, with a strong bias toward inclusion in either neurons or muscle cells (**Supplemental Table S3**; Koterniak et al, in preparation). The PAM scores for our 20 PRA-identified ISE and ISS k-mer motifs were calculated for 5′ and 3′ orthologous intronic regions flanking these switch-like exons from up to 50 *Caenorhabditis* species (Fusca et al., 2025). We then performed hierarchical clustering on PRA ISE/ISS motifs and intronic fragments based on PAM scores results to identify potential co-regulated exons (**Fig. 6A**).

**Figure 6:**
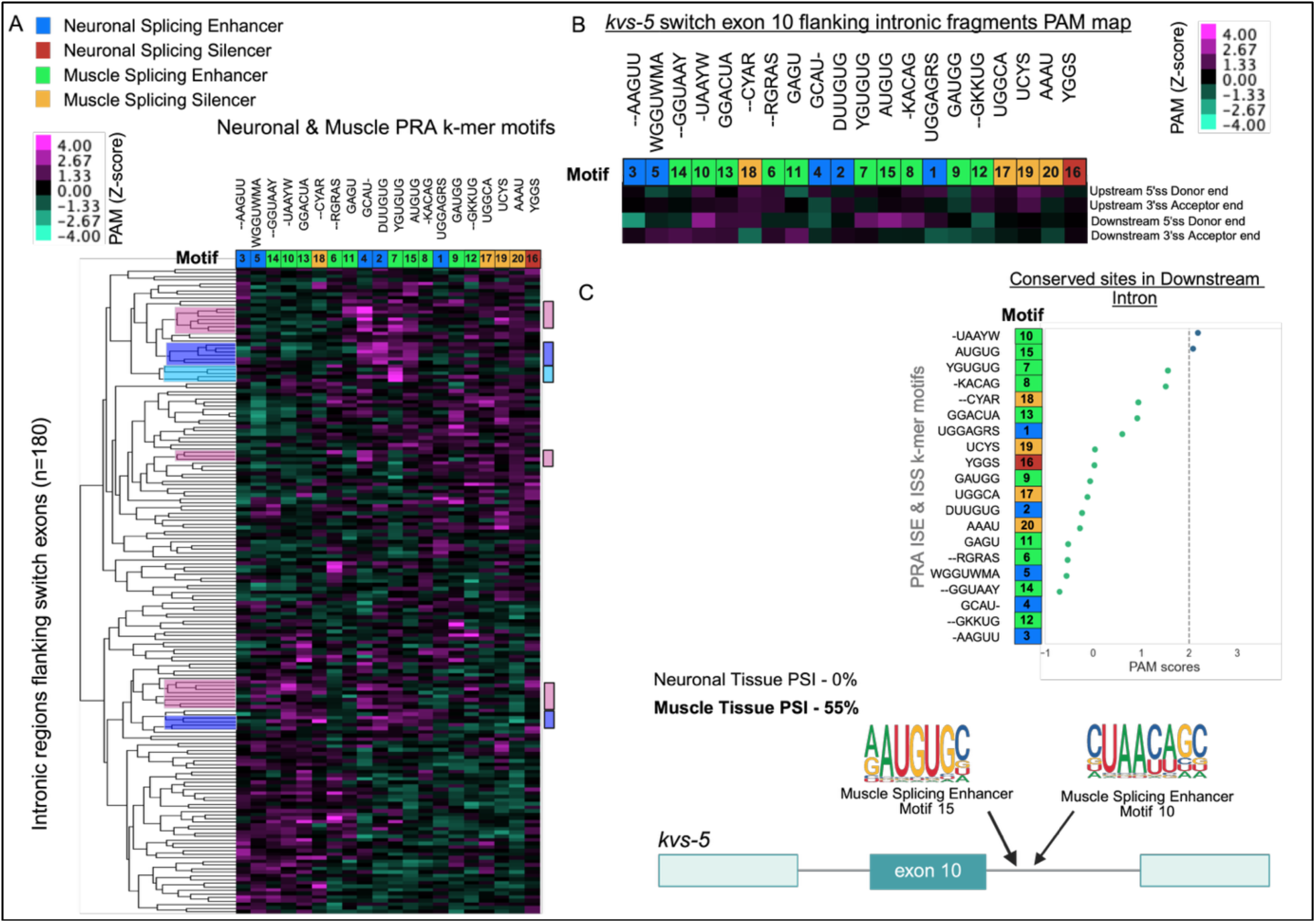
PRA-derived ISE and ISS motifs can be mapped to endogenous conserved intronic regions flanking tissue-biased switch-like exons. **(A)** PAM Heatmap showing conservation patterns of PRA splicing motifs across 180 intronic fragments extracted from 47 *C. elegans* tissue-differentially spliced genes. The hierarchical clustering of intronic fragments (Y-axis) and PRA ISE and ISS k-mer motifs (X-axis) based on PAM scores results in the grouping of intronic fragments containing similar conserved PRA motif patterns. The legends on the left show PAM scores and the colours used for different PRA ISE and ISS categories; the tissue-specific PRA ISE and ISS k-mer motifs are indicated accordingly on the X-axis of the heatmap along with their consensus sequence. Most prominent clusters include neuronal splicing enhancers motif 4 GCAU- (intronic fragment group highlighted in pink), motif 2 DUUGUG and muscle splicing enhancer motif 7 YGUGUG. The intronic fragment groups with elevated PAM scores for these motifs are highlighted in pink, light purple and sky-blue colors. **(B and C)** Focusing on a specific example, *kvs-5*, the intronic fragments PAM map with 5’ss and 3’ss containing regions within introns adjacent to the switch-like exon in *kvs-5* is presented. It shows the presence of conserved muscle-biased splicing-enhancing motifs identified from the PRA screen, in the downstream intron of the *kvs-5* tenth exon. The tenth exon undergoes splicing inclusion in a muscle-tissue biased manner (muscle PSI 55% compared to neuronal PSI of 0%; based on tissue-specific profiling data (Koterniak et al. 2020)).

Among the 20 PRA-identified tissue-biased ISE and ISS k-mer motifs, 19 were found to be conserved in intronic regions adjacent to switch-like exons and the orthologous sequences of these intronic regions in other *Caenorhabditis* species with a PAM score > 2 (**Fig. 6A**). As expected, the neuron-biased ISE motif 4 (--GCAUG--), resembling the FOX-1/ASD-1 motif (**Fig. 6A**), was found highly conserved (PAM score ≥ 2) in 20 intronic regions from 18 splicing events. Similarly, neuron-biased ISE motif 2 (-UUGUG) that resembles the core UNC-75 motif (Supplemental Fig. S3), and muscle-biased ISE motif 7 (YGUGUG) that closely matches the SUP- 12 motif, were found conserved in 9 and 8 intronic regions, respectively **(Fig. 6A)**.

Interestingly, in addition to introns containing single highly conserved motifs, a subset (20%) involved two or more conserved k-mer motifs, suggesting combinatorial control (**Fig. 6A**). Focusing on one example involving robust muscle-specific exon inclusion, we present the PAM map for the intronic regions flanking alternative exon 10 in the *kvs-5* gene, which codes for a voltage-sensitive channel subunit (Meissner et al., 2011) (Fig. 6B and 6C; neuronal PSI = 0% and muscle PSI 55%; **Supplemental Table S3; Fig. 6B and C**). Notably, the downstream intron of this exon consists of highly conserved muscle-biased ISE motifs (cluster 10 and cluster 5). The presence of these motifs with muscle-biased ISE activity at relevant intronic locations suggests that they may serve a key role in promoting muscle-biased inclusion of alternative exon 10 in this gene (**Fig. 6B and 6C**). Taken together, our results demonstrate that ISE and ISS motifs identified by our PRA analysis are conserved across nematodes in intronic regions flanking tissue-biased alternative exons, consistent with a role in alternative splicing regulation of endogenous transcripts. Moreover, our data suggest that the splicing of these switch-like exons can be regulated through diverse combinations of *cis* elements. Finally, by combining PAM score representations, PRA activity measurements, and empirically determined splicing outcomes, testable predictions can be made about how the presence of specific *cis* elements can lead to tissue-biased splicing patterns.

### Mutation of PRA-identified ISEs disrupts the splicing of an endogenous neuronal microexon

To further demonstrate the activity of the PRA-identified ISE and ISS motifs, we performed targeted mutagenesis experiments in the context of a native tissue-biased alternative exon. The *dpy-23* gene is known to enable clathrin adaptor activity, playing a crucial role in trafficking and endocytosis (Hermann et al., 2005; C. L. Pan et al., 2008). Interestingly, the fourth exon in the *dpy-23* gene is a microexon (18nt) that undergoes pronounced neuron-biased inclusion (neuronal PSI = 82% and muscle PSI 10%) (Koterniak et al., 2020). The PAM signatures for the intronic regions flanking the microexon revealed that two motifs with neuron-biased ISE activity (motif 2 and motif 3) were found to be conserved in the downstream intron towards the 5′ splice site end **(Fig. 7A and 7B)**. Thus, we hypothesized that these *cis* elements may play a key role in promoting inclusion of this microexon in neurons.

**Figure 7:**
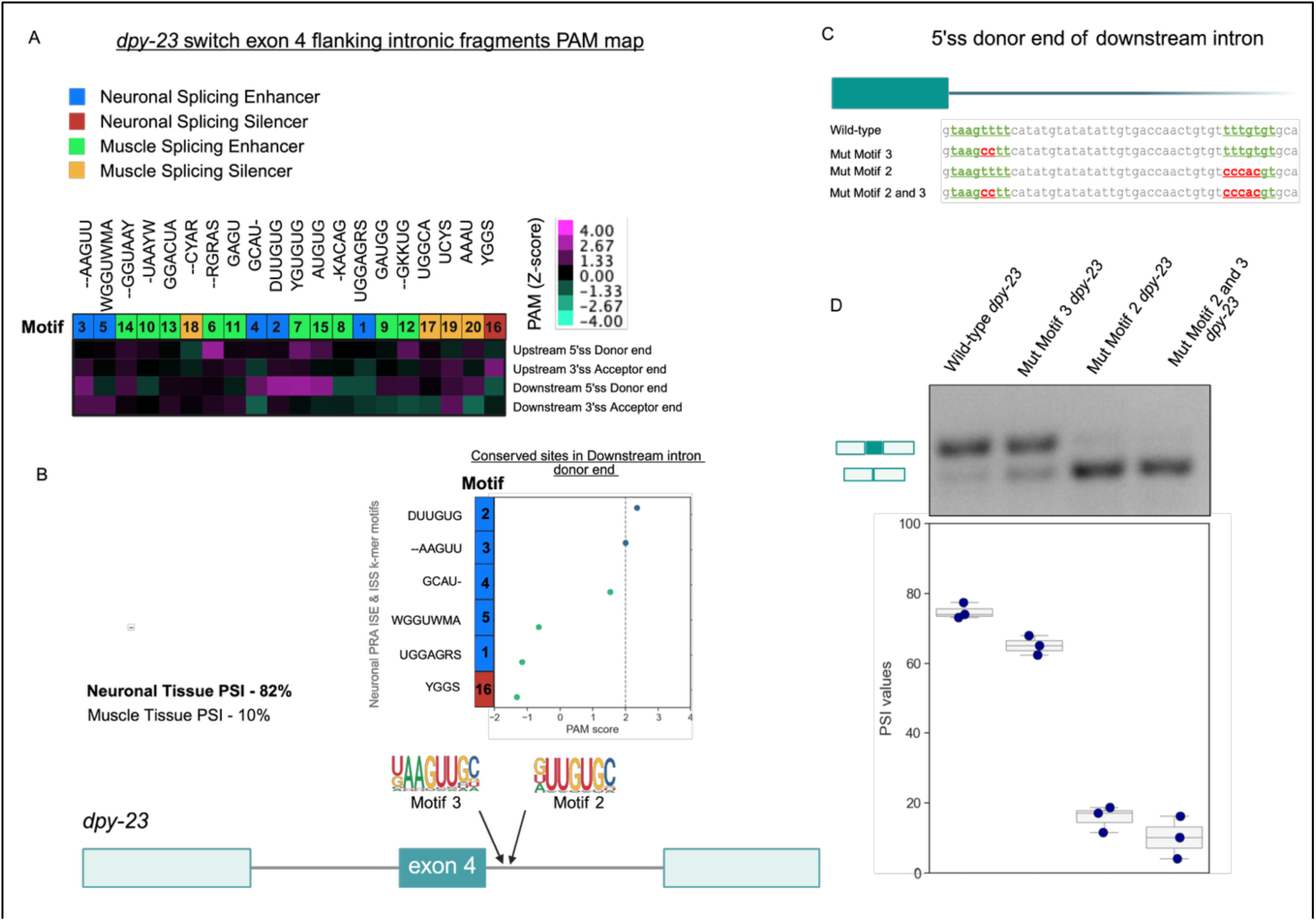
Identification and mutagenesis of neuron-biased PRA ISE k-mers found conserved in intronic regions flanking tissue-differentially spliced microexon 4 in the *dpy-23* gene **(A)** Heatmap shows PAM scores for neuron- and muscle-biased ISE and ISS motifs adjacent to the tissue-differentially spliced microexon 4 in *dpy-23*. The legend on the right shows the PAM scores and the legend on the left shows colors used for different PRA ISE and ISS k-mer motif categories as shown on the X-axis of the heatmap. **(B)** *dpy-23* microexon 4 is differentially spliced between neuronal and muscle tissue. It is highly included in neuronal tissue (82%) compared to muscle tissue (10%). As shown in the schematic, neuronal splicing enhancers (NeuSE) motif 2 and motif 3 (PAM score ≥ 2) were found conserved in the donor end region of downstream intron. **(C)** The downstream intronic sequence is shown with NeuSE motif 3 adjacent to the 5′ splice site and NeuSE motif 2 located 32 bp downstream. Three mutant minigene constructs were generated: one with motif 3 mutated (Mut Motif 3), one with motif 2 mutated (Mut Motif 2), and one with both sites mutated (Mut Motif 2 and 3). **(D)** Semi-quantitative RT-PCR, coupled with densitometry, was used to evaluate splicing outcomes in *C. elegans* expressing wild-type and mutant *dpy-23* minigenes.

To test this hypothesis, we introduced site-specific mutations within the mapped motifs, either alone, or in combination, in a *dpy-23* minigene splicing reporter that recapitulates tissue-biased microexon inclusion **(Fig. 7C).** Notably, all neuron-expressed *dpy-23* minigene reporter plasmids with mutated PRA splicing motifs tested exhibited reduced microexon inclusion **(Fig. 7D)**. The reporter plasmid with mutated motif 3 exhibited a moderate decrease in microexon inclusion with the average PSI value dropping from 74.86 +/- 2.2% for the wild-type neuronal *dpy- 23* minigene reporter to 65.14 +/- 2.7%. In contrast, mutation of motif 2 resulted in a substantially stronger effect (PSI = 15.88 +/- 3.7%, mean dPSI compared with wild-type = -59%; **Fig.7D**), highlighting a critical role of this motif in *dpy-23* microexon splicing regulation. The reporter with both motif 2 and motif 3 mutated exhibited a further reduction in microexon inclusion (PSI = 10.14 +/- 6%). However, effects were comparable to that observed with mutation of motif 2, suggesting a dominant role of this motif in *dpy-23* microexon splicing regulation **(Fig. 7D)**. Collectively, these results demonstrate that intronic elements identified in our PRA screen display tissue-biased splicing regulatory activity *in vivo*.

## Discussion

### A screening tool to identify splicing regulatory sequences with tissue-biased activity in vivo

In this study we have successfully implemented an *in vivo* PRA to investigate the splicing regulatory effects of thousands of synthetic intronic sequences in a quantifiable manner *in vivo*. We further demonstrate that by adapting this methodology for a multicellular animal, *cis* elements with tissue-biased activity can be extracted from the data. Our approach thus enables the interrogation of splicing regulatory signals in their physiologically relevant cellular environment, rather than in cell lines, complementing ongoing PRA-based efforts to study *cis* regulatory logic of gene expression in live organisms (Brown et al., 2025; Lambert et al., 2021; Zheng et al., 2023). We anticipate that our *in vivo* PRA approach can be expanded as a screening tool to identify splicing regulatory sequences with diverse spatio-temporal activity in the context of animal development. Moreover, given the wealth of environmental stimuli and signals known to impact gene expression in *C. elegans* (Lee et al., 2001; Martinez et al., 2020; Tamburino et al., 2013), our PRA will also prove useful to identify intronic sequences that show signal-dependent splicing activity in the context of these paradigms.

The intronic splicing enhancer and silencer elements we have identified provide a subset of the parts list contributing to how context-dependent splicing is achieved. Our work thus complements ongoing efforts to develop interpretable models for the rules governing splicing outcomes across organisms, also known as the splicing code (K. Gupta et al., 2024; La Fleur et al., 2024; Liao et al., 2023). Among the sequences we found with regulatory activity are those recognized by RNA binding proteins with known tissue-biased impacts on alternative splicing (Jin et al., 2003; Norris et al., 2014; J. H. Tan & Fraser, 2017). However, we also discovered sequences with strong effects on splicing without clear cognate RBP partners (‘orphan elements’). Future work will be directed to deorphanize these *cis* elements through biochemical or reverse genetic screening approaches, identifying novel regulators of splicing in native tissue contexts.

In addition to identifying intronic enhancers and silencers of alternative splicing, our screen identified candidate 5′ splice sites that effectively competed with the naturally occurring 5′ splice site flanking the reporter alternative exon (**Fig. 2**). Consistent with early studies (Eperon et al., 1986) we identify that a GUAAG pentamer, which is perfectly complementary to nucleotides in the U1 snRNA, is often sufficient to induce high levels of cryptic splicing at the expense of the nearby native alternative splice site. However, a somewhat more surprising result was that even non-canonical splice sites, including those with GC at positions +1 and +2 (GC-splice donors), led to intermediate levels of cryptic splicing when positioned downstream of the native alternative splice donor. Consistent with this result, a small proportion of introns in *C. elegans* and other metazoa contain GC-splice donors associated with both constitutive and alternative spliced exons (Farrer et al., 2002). Additionally, our results support growing evidence that effective splice donor sequences can exhibit more flexibility in sequence composition than previously appreciated, creating additional opportunities for context-dependent splicing regulation when combined with other enhancer and silencer motifs (Roca et al., 2003, 2012; J. Tan et al., 2016; Wong et al., 2018; Yu et al., 2008). Finally, our results also predict that identified cryptic splice-site donor sequences should be selected against and depleted from introns, particularly those flanking alternative exons with weakened 5′ splice sites.

With the rise of single cell RNA-seq (scRNA-seq) approaches in *C. elegans* (Barrett et al., 2025; Cao et al., 2017; Packer et al., 2019; Roux et al., 2023; Taylor et al., 2021), future iterations of our *in vivo* PRA can also be modified to survey the effects of randomized sequences on splicing outcomes in single cells. Indeed, such methods coupling *in vivo* PRAs with scRNA-seq are being developed for other layers of gene expression (Zhang et al., 2025; Zhao et al., 2023). We note that at present, especially for the nervous system, which contains diverse neuronal subtypes, our broad tissue *in vivo* PRA is likely enriching for signals that act broadly in neurons. However, analogous to vertebrates, recent studies have shown alternative splicing differences between individual neuronal subtypes (Feng et al., 2021; Thompson et al., 2019; Weinreb et al., 2025; Wolfe et al., 2025). We thus anticipate that sampling the activity of randomized sequences at single cell resolution will be of relevance to the nervous system.

### Extensive combinatorial control of alternative splicing supported by evolutionary signatures

Mapping our *in vivo* PRA-derived *cis* elements onto native intronic sequences that flank switch-like exons revealed evolutionary-conserved signals for these motifs (Fig. 6). Our work builds upon previous studies demonstrating that comparative genomics can be useful to identify splicing regulatory sequences in intronic regions (Bortfeldt et al., 2008; Kabat et al., 2006; Kol et al., 2005; Yeo et al., 2007). Perhaps most striking, although clustering identified several shared intronic motif signatures, we often observed that diverse and unique combinations of *cis* elements are enriched within individual introns.

These results have several broad implications. First, the data indicate that there are many paths to achieve tissue-biased splicing outcomes, as indicated by the growing repertoire of nuclear RBPs in multicellular animals, including *C. elegans*, that can serve this role (Barberan-Soler et al., 2011; Koterniak et al., 2020; Peng & Murray, 2025; Tan & Fraser, 2017; Laver et al., biorxiv). Such flexibility would accommodate the gain and loss of individual sequences through evolution, supporting compensatory mechanisms to preserve regulatory function (similar to ‘developmental systems drift’; True & Haag, 2001), or contributing to novel splicing patterns (Barbosa-Morais et al., 2012; Jelen et al., 2007). Second, the utilization of several distinct *cis* elements would also enable multiple RBPs to work additively, antagonistically, or synergistically, to create robust splicing outcomes during development and between tissues, which we and others have observed *in vivo* (Dvinge, 2018; Iadevaia & Gerber, 2015; Jangi & Sharp, 2014; Lin & Tarn, 2005; Norris et al., 2014; J. H. Tan & Fraser, 2017; Ule & Blencowe, 2019). Finally, these results are reminiscent of ‘phenotypic convergence’, where within a given promoter or intronic region, distinct *cis* elements are recognized by their corresponding trans-acting factors in different cell types, to achieve the same regulatory outcome (Carrasco et al., 2020; Kuroyanagi et al., 2006). Our study thus provides a framework to integrate PRA screens with comparative genomics to make informative and testable mechanistic predictions.

### Two intronic enhancers regulate the inclusion of a neuronal microexon in C. elegans

Guided by our PRA and PAM score analysis, we validated the impact of two neuron-biased intronic splicing enhancers on the splicing of exon 4, an 18 nucleotide microexon, from the *dpy- 23* gene. Microexons (less than 27 nucleotides) have emerged as an interesting class of exons, with an increased frequency in vertebrate genes compared to invertebrates. Aberrant regulation of these small exons has been associated with autism spectrum disorders and pancreatic cancer (Gonatopoulos-Pournatzis & Blencowe, 2020; Irimia et al., 2014). The increased occurrence of microexons in vertebrates is thought to be associated with the emergence of the SR-related splicing activator proteins SRRM3 and SRRM4 (Head et al., 2021; Irimia et al., 2014). However, perturbation of SRRM4 does not eliminate inclusion of all microexons, and other factors have also been found to work together with this protein (T. Gupta et al., 2025). *C. elegans* lacks an SRRM4 ortholog, but its genome does contain microexons which can be spliced in a tissue-biased manner (Choudhary et al., 2021; Torres-Méndez et al., 2019) . Interestingly, the intronic splicing enhancer that we validated to have a strong impact on *dpy-23* microexon splicing matches the core UUGUG consensus sequence recognized by the CELF ortholog UNC-75 (Norris et al., 2014; Pilaka-Akella et al., 2025). This result is consistent with previous studies suggesting an evolutionary conserved role for CELF proteins in regulating microexon splicing in planaria and in vertebrates (Solana et al., 2016). In *C. elegans*, we add UNC-75/CELF to an emerging list of conserved intron-binding RBPs that likely work combinatorially with PRP-40 (a U1 snRNP complex-interacting protein) to regulate microexon inclusion (Choudhary et al., 2021).

The second neuron-biased intronic splicing enhancer we identified had a subtle but significant effect on microexon splicing (Fig. 6). At present, it is unclear what factor may be recognizing the core AAGUU sequence in this enhancer, although a likely candidate could be the Hu ortholog EXC-7, which recognizes a similar motif (Norris et al., 2014). Consistent with this possibility, EXC-7 has been found to regulate a subset of microexons in *C. elegans* (Choudhary et al., 2021) and is also known to co-regulate alternative splicing with UNC-75 (Norris et al., 2014). The proximity of this enhancer to the 5′ splice site flanking the microexon may create an environment that favours tissue-biased inclusion in neurons, possibly through stimulating or stabilizing U1 snRNP recruitment, analogous to the TIA1/TIAL proteins (Förch et al., 2002).

Further exploration of this enhancer will shed additional mechanistic insight into its role in tissue-biased splicing. Collectively, these results highlight the broader utility of intersecting our PRA- derived regulatory sequences with endogenous alternatively spliced loci to inform novel regulatory landscapes. We anticipate that continued use of these screening approaches will prove useful in our efforts to understand the code governing context-specific splicing in multicellular animals.

## Data Availability

All code and raw sequence data (fastq files) are available at: https://github.com/sanjanabhatnagar/in-vivoPRA-and-C.elegans-PAM-paper.git https://github.com/sanjanabhatnagar/Inferring-Exon-and-Intron-Metadata-from-.gff-file.git

## Supplementary Data Statement

Supplementary Data are available at *NAR* Online.

## Acknowledgments

We thank members of the Calarco and Moses labs for critical discussions throughout the course of this study. We also acknowledge Santiago Vargas and Alissa Hofmann for early work on this project.

## Author Contributions Statement

Conceptualization: S.B., A.M.M., J.A.C.; Data curation: S.B, M.Z, B.K., D.F.; Formal analysis: S.B., N.H.S, M.Z, B.K., D.F; Funding acquisition: A.M.M, J.A.C; Investigation: S.B, J.H., N.H.S., M.Z, Y.L., I.S., B.K., D.F.; Methodology: S.B., A.M.M, J.A.C.; Project administration: S.B., J.A.C.; Resources: A.D.C., A.M.M., J.A.C.; Software: S.B., A.M.M.; Supervision: A.D.C., A.M.M., J.A.C.; Validation: S.B., N.H.S., I.S.; Visualization - S.B., J.A.C.; Writing – original draft: S.B., J.A.C.; Writing – review & editing: S.B., D.F., A.D.C., A.M.M., J.A.C.

---

**Supplementary Figure S1:**
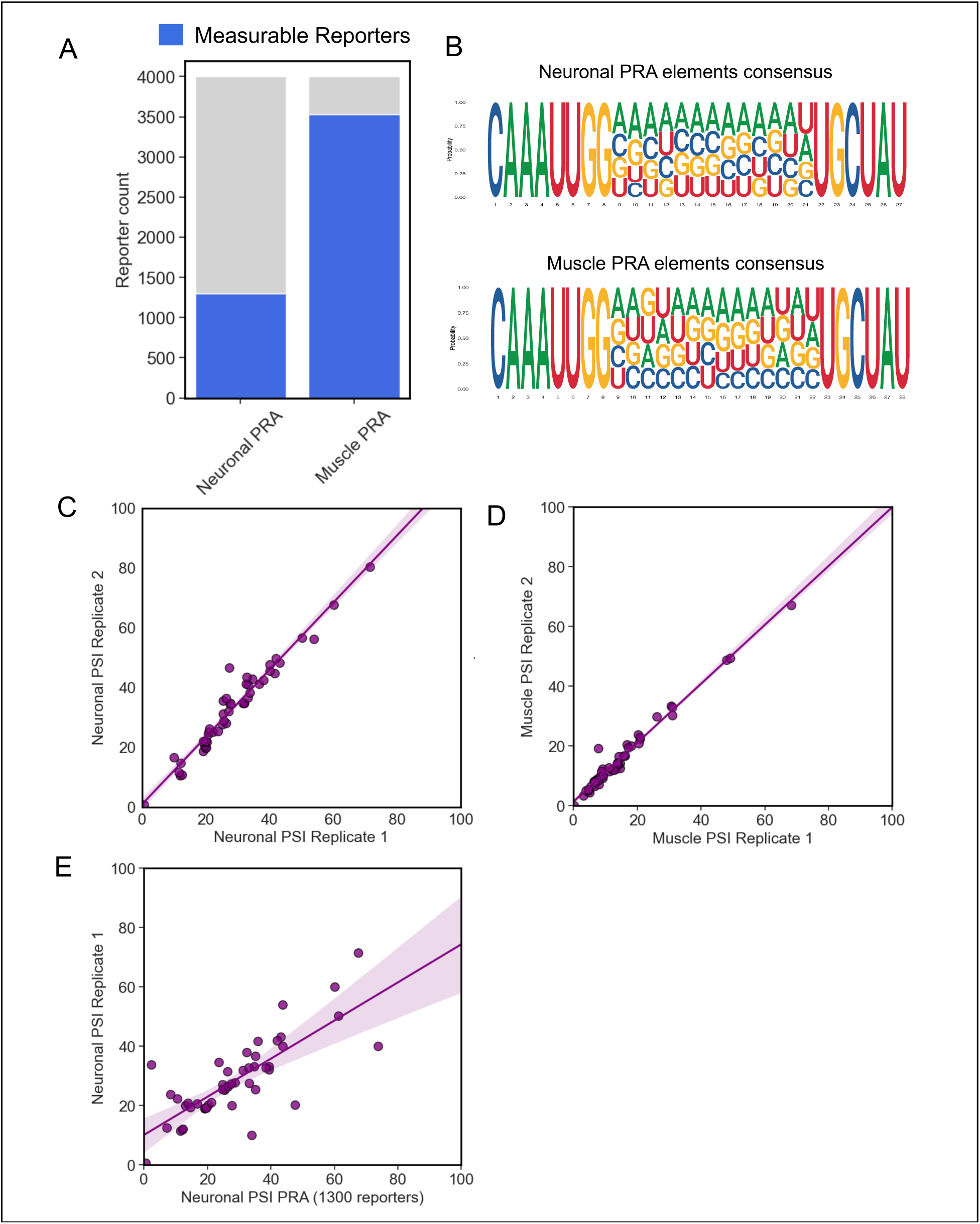
Validation of robustness and randomization in neuronal and muscle PRA **(A)** From 4000 reporter neuronal and muscle PRA libraries were injected into *C. elegans*, after the analysis of PRA RNA-seq data, transcripts were mapped back to 1337 neuronal PRA reporters and 3642 muscle PRA reporters. These reporter transcripts were used for downstream analysis. **(B)** Consensus sequences generated by multiple sequence alignment of introduced 10bp PRA intronic elements confirm randomization in both tissue contexts. **(C-E)** The scatter plots show a high positive correlation between PSI value measurements of 71 reporters from smaller-scale neuronal and muscle PRA replicates and neuronal smaller-scale PRA vs. neuronal 4000 reporter PRA, thus validating the robustness of the screen.

**Supplementary Figure S2:**
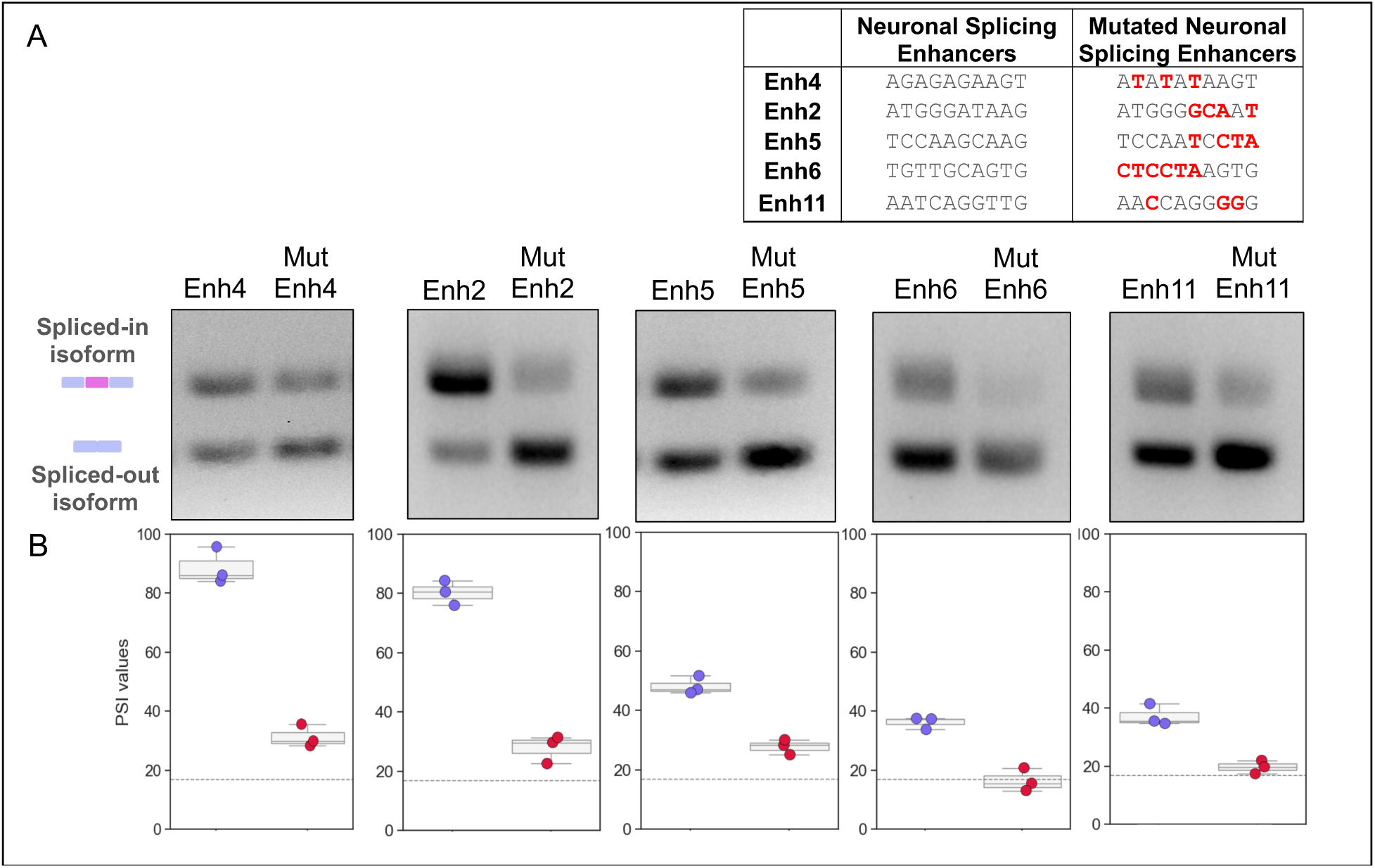
Random mutagenesis of neuronal PRA splicing enhancing intronic elements results in loss of splicing enhancement activity **(A)** Five neuronal PRA ISEs were selected for random mutagenesis, followed by semi-quantitative RT-PCR to observe the altered splicing patterns. Two bands were obtained for all reporters: the spliced-in isoform and the spliced-out isoform. The sequences of the elements in the PRA reporters and the mutated PRA reporters that were injected individually into N2 adults are shown in the table on the right side. **(B)** Gel band intensities were quantified, and densitometry-based PSI values (PSI_densitometry_) were calculated across three replicates and compared. The splicing patterns of the reference reporter is shown using the gridline, (PSIRNA-seq of reference reporter is 17%).

**Supplemental Figure S3:**
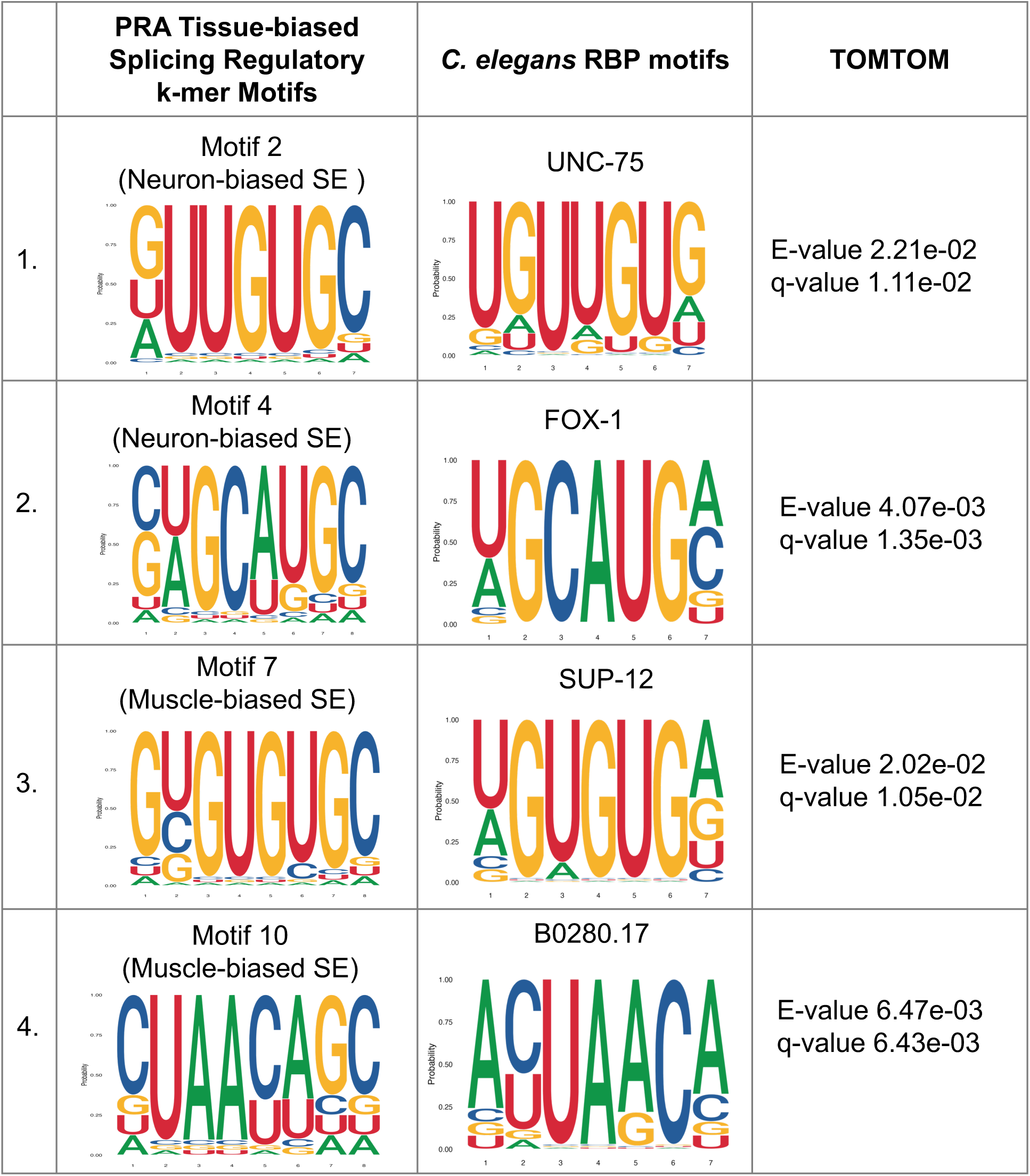
Comparative analysis of muscle and neuronal PRA ISS and ISE k- mer clusters and 38 C. elegans RBP motifs. TOMTOM was used for motif comparison analysis (See Methods) of PRA ISE and ISS k-mer motifs and known *C. elegans* RBP motifs. The last column shows the confidence scores for the corresponding matches.

